# Endothelial-secreted Endocan protein acts as a PDGFR alpha ligand and regulates vascularity, radioresistance, and regional phenotype in glioblastoma

**DOI:** 10.1101/2020.10.12.335091

**Authors:** Soniya Bastola, Marat S. Pavlyukov, Yasmin Ghochani, Neel Sharma, Mayu A. Nakano, Sree Deepthi Muthukrishnan, Sang Yul Yu, Min Soo Kim, Alireza Sohrabi, Natalia P. Biscola, Daisuke Yamashita, Ksenia S. Anufrieva, Riki Kawaguchi, Yue Qin, Stephanie K. Seidlits, Alma L Burlingame, Juan A. Oses-Prieto, Leif A. Havton, Steven A. Goldman, Anita B. Hjelmeland, Ichiro Nakano, Harley I. Kornblum

## Abstract

One of the hallmarks of glioblastoma (GBM) is extensive neovascularization. In addition to supplying blood and nutrients, vascular endothelial (VE) cells provide trophic support to GBM cells via paracrine signaling, the precise mechanisms of which are being unraveled. Here, using patient-derived GBM and VE cells as well as orthotopic GBM mouse models, we report that Endocan (*ESM1*), an endothelial-secreted proteoglycan, confers enhanced proliferative, migratory, and angiogenic properties to GBM cells and regulates their spatial identity. Mechanistically, Endocan exerts at least part of its functions via direct binding and activation of the PDGFRA receptor. Subsequent downstream signaling enhances chromatin accessibility of the Myc promoter and upregulates Myc expression inducing highly stable phenotypic changes in GBM cells. Furthermore, Endocan confers a radioprotection phenotype in GBM cells, both *in vitro* and *in vivo*. Inhibition of Endocan-PDGFRA signaling with ponatinib increases survival in the *Esm1* wild-type but not in the *Esm1* knock-out mouse GBM model. Our findings identify Endocan and its downstream signaling axis as a potential target to subdue the recurrence of GBM and further highlight the importance of vascular to tumor cell signaling for GBM biology.

**Significance statement:** Identification of the Endocan/PDGFRA/Myc axis demonstrates an important role of VE cells in GBM malignancy. The contribution of Endocan to the development of GBM cell populations with different phenotypes reveal an additional pathway underlying the origin of GBM intratumoral heterogeneity. Targeting Endocan-mediated crosstalk may enhance the efficacy of GBM treatment.

## Introduction

Glioblastoma (GBM) is a highly aggressive, vascular-rich tumor that exploits vascular endothelial (VE) cells to contribute to its growth^1,2^. VE cells provide paracrine factors, nutrition, oxygen, as well as a niche for glioma stem cells (GSCs)^1,3,4^. GSCs residing within the perivascular niche maintain the capacity for self-renewal and promote GBM progression, and recurrence^3,5^. Developing an understanding of the basic biology of tumor-associated vasculature will open the door to new therapeutic targets and the establishment of novel therapies.

In experimental models, tumor-associated VE cells support the maintenance of GSCs^3^. This function can be recapitulated by VE cell-derived conditioned media (CM), suggesting that VE cells produce soluble factors that influence the GSC phenotype. In turn, GSCs can secrete angiogenic factors to promote further neovacularisation^6^. Our recent work has established an interactome network within GBM-associated VE cells and identified key dysregulated genes encoding secreted proteins emanating from VE cells^7^. One such gene is *ESM1* that encodes Endocan, a soluble proteoglycan of 50 kDa, constituted of a mature polypeptide of 165 amino acids (18 kDa) and a single dermatan sulphate chain covalently linked to the serine residue at position 137^8^. *ESM1* expression is regulated by the VEGF/hypoxia pathway and serves as a marker of endothelial cell activation during neoangiogenesis in renal, non-small cell lung, hepatocellular, bladder carcinomas, and GBM^8–10^. *ESM1* is significantly upregulated in tumor-derived vascular cells compared to non-transformed VE cells and its level correlates with higher grade and shorter survival in glioma^11,12^. However, the functional role of Endocan in glioblastoma and, subsequently, its potential as a therapeutic target remains unknown.

In this study using patient-derived GBM and VE cells, syngeneic orthotopic GBM models, and an *Esm1* knockout mouse model^13^, we investigated the role of Endocan in intercellular crosstalk between GBM and VE cells. Our findings suggest that targeting Endocan itself or downstream signaling pathways activated by this protein may represent a novel therapeutic approach in the treatment of GBM.

## Results

### Vascular-secreted Endocan promotes proliferation and migration of GBM cells

To understand how VE cells contribute to the progression of GBM, we tested whether VE cells promote the aggressiveness of GBM *in vitro* and *in vivo* by using two VE cell models: a short-term primary culture of glioblastoma patient-derived VE cells (TEC15) and an immortalized brain VE cell line HBEC-5i. First, we demonstrated that conditioned media (CM) collected from both TEC15 and HBEC-5i cells promote *in vitro* proliferation of gliomasphere lines obtained from four different patients **(Figure 1a** and **Figure S1a)**. Next, we co-injected VE cells (TEC15 or HBEC-5i) with the patient-derived gliomasphere line 1051^14^ at a 1:10 ratio into SCID mice, and showed that they significantly shortened animal survival **(Figures 1b, S1b)**. To identify potential trophic factors that could be elaborated by tumor-associated VE cells, we previously isolated CD31^+^ tumor-associated vascular cells from 8 glioblastoma cases along with non-cancerous CD31^+^ cells from 5 craniotomies for epilepsy as controls^7^. Subsequent RNA-sequencing (RNA-seq) revealed *ESM1* as the most significantly upregulated gene in tumor-derived CD31^+^ cells compared to the non-neoplastic counterparts (**Figure 1c)**. Analysis of a previously published single cell RNAseq database^15^ confirmed that *ESM1* is upregulated in one of the endothelial sub-clusters, which was characterized by gene signatures associated with endothelial tip cell formation, tumor angiogenesis, and vascular basement membrane remodeling (**Figures S1c,d**). In the TCGA database, *ESM1* expression was found to correlate with higher glioma grade (**Figure S1e**) and shorter survival of GBM patients (**Figure S1f**). ELISA analysis demonstrated that Endocan, the *ESM1* gene product, was secreted by HBEC-5i and glioblastoma patient-derived VE cells (TEC 14 and TEC15) at concentrations exceeding 100pg/ml, while conditioned media (CM) from normal human astrocytes (NHA) and gliomasphere lines (1079, 157, 711, and 1051) contained nearly undetectable amounts of Endocan (**Figure 1d**). We further confirmed this result by immunohistochemical staining (IHC) and demonstrated that Endocan is exclusively co-localized with CD31+ VE cells in the perivascular areas of our GBM clinical samples (**Figure 1e**).

**Figure 1:**
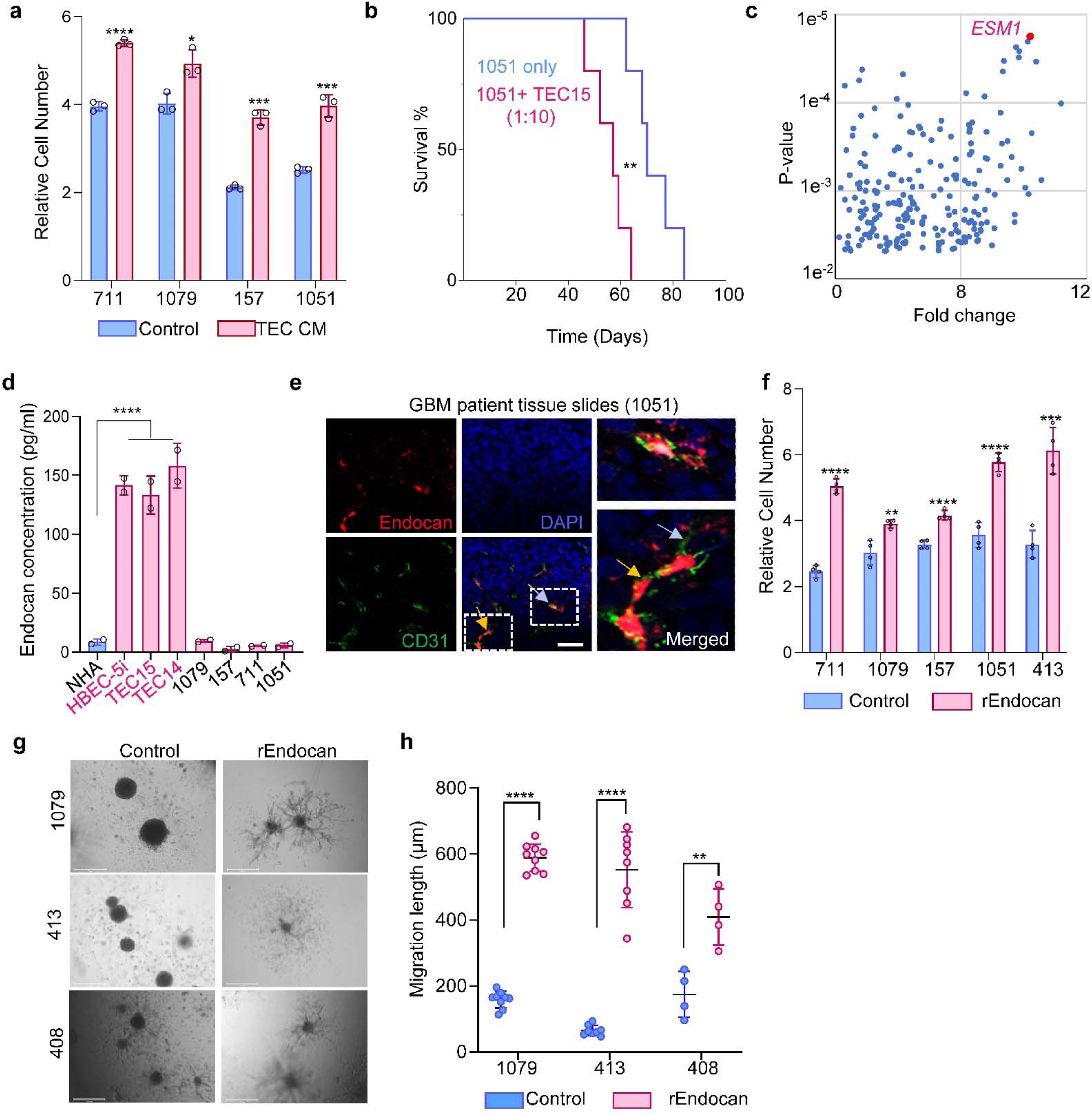
Vascular-secreted Endocan promotes proliferation and migration of GBM cells. **A,** Proliferation of GBM cells (711, 1079, 157, 1051) treated with control media or conditioned media collected from Tumor Endothelial Cells (TEC CM); n=3 independent experiments. \**P=0.015, **P=0.007*, *\*\*\*P=0.0001*, and *\*\*\*\*P<0.0001*. **D**, ELISA assay comparing levels of secreted Endocan in CM from Normal Human Astrocytes (NHA), Human Brain Endothelial Cells (HBEC-5i), Tumor-associated Endothelial cells (TEC15, TEC14), and GBM cells (1079, 157, 711, and 1051). One-way Anova and Tukey post-hoc test, n=2 independent experiments, *\*\*\*\*P<0.0001*. **E**, Representative immunofluorescence (IF) staining for Endocan (red) and CD31 (green) in formalin-fixed paraffin-embedded tissue sections from glioblastoma patient 1051. Nuclei stained with DAPI (blue). Scale bar, 50 μm. **F**, Proliferation of GBM cells (711, 1079, 157, 1051, 413) with or without rEndocan (10ng/ml). Proliferation was measured 5 days after treatment; n=4 independent experiments; *\*\*P<0.005, ***P<0.0005, ****P<0.0001*. **G**, Representative microscopic images of control and rEndocan treated (10ng/ml) GBM spheroids encapsulated in the hydrogel. Images were taken 3 days post encapsulation. n=6 spheroids per group. Scale bar, 500µm. **H,** Quantification of migration distance in control and rEndocan spheroids formed by 408, 413 or 1079 cells. Data from n= 4-10 spheroids were used for analysis. \*\**P*=0.0053; \*\*\*\**P*<0.0001. All quantitative data are average ±SD, unpaired two-tailed Student’s t test was used to determine the significance of differences between the indicated groups where applicable.

Next, we investigated whether Endocan could influence GBM phenotype. Incubation with recombinant Endocan (rEndocan) promoted the proliferation of multiple GBM sphere lines **(Figure 1f)**. As GBM cells are known to migrate along blood vessels^16,17^, we also examined the effects of Endocan on GBM cell migration using a previously described hydrogel-based system^18^. rEndocan-treated spheroids derived from 3 different patients had significantly enhanced motility as revealed by both migration length (**Figures 1g,h**) and shape factor analysis (**Figure S1g**).

Collectively, these findings suggest that vascular-secreted Endocan drive both GBM cell proliferation and migration.

### Endocan is essential for establishing the hypervascular phenotype of GBM

To address the potential role of paracrine Endocan signaling in crafting the GBM phenotype *in vivo*, we utilized wild-type (WT) and *Esm1* knockout (*Esm1* KO) mice^13^ as host systems for implanting glioma cells derived from two different murine GBM models. First, we used *in vitro* gliomasphere cultures termed as 7080^19^ that was derived from mice harboring mutations in *p53*, *Pten*, and *Nf1* and enriched for the tumor initiating subpopulation^20^. Second, we utilized freshly-resected murine glioblastoma-like cells isolated from tumors induced by RCAS-PDGAB lentivirus injection into Nestin-Tva/Cdkn2a-/- mice^21^. Both types of cancer cells were injected into the brains of WT and *Esm1* KO mice. The resultant tumors in WT mice exhibited numerous intratumoral hemorrhages in sharp contrast to grayish necrotic tumors in *Esm1* KO mice (**Figures 2a, S2a**). Subsequent IHC for CD31 demonstrated significantly fewer CD31^+^ VE cells in the tumors from *Esm1* KO animals with reduced vessel diameter as compared to those formed in wildtype animals (**Figures 2a-b**, **S2a-b**). At the ultrastructural level, 7080 tumors in the WT host exhibited proliferative blood vessels with hypertrophy and the basement membrane showed an irregular outline and varying diameters, typical of human glioblastoma^16,22^, while in the *Esm1* KO host tumors exhibited sparse blood vessels with a regular basement membrane measured as Blood vessel ratio-SAE using previously defined methods^23^ (**Figures 2c-d**).

**Figure 2:**
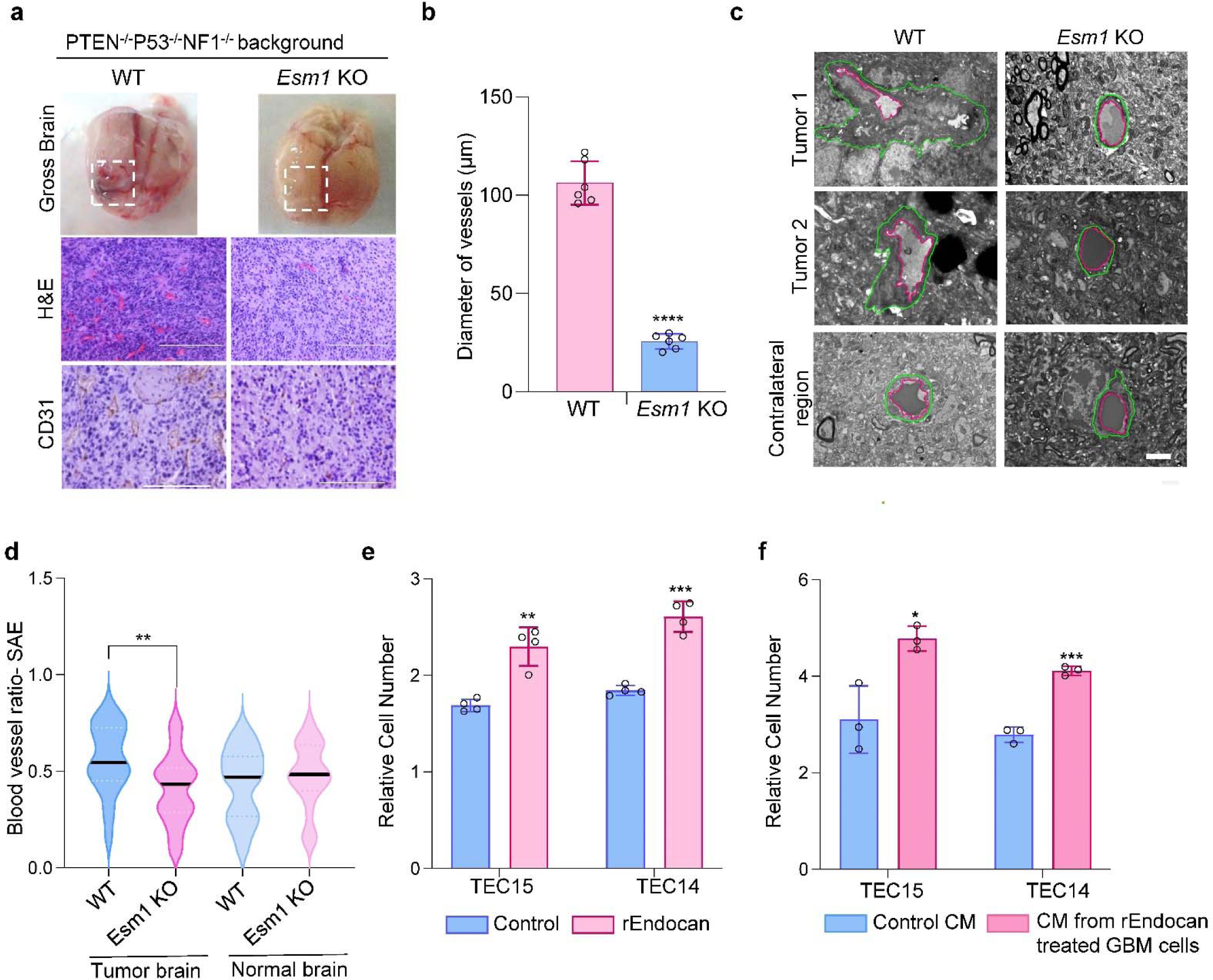
Endocan is essential for establishing the hypervascular phenotype of GBM. **A,** Images of the brains collected from 7080 glioma-bearing *Esm*1 WT or *Esm1* KO mice (upper panel). H&E staining of the tumor slices (middle panel, Scale bar, 200 μm) and CD31 staining of the tumor slices (lower panel, Scale bar, 200 μm). **B**, Quantitation of the diameter of the blood vessels formed in tumors from *PTEN/P53/NF1* deleted glioblastoma model. CD31 positive VE cells were imaged and then measured in ImageJ using ImageJ vessel analysis tool. n=5 mice per group, *\*\*\*P<0.0001*. **C**, Representative transmission electron microscopy images of tumors formed in *Esm1* WT and KO mice (middle panel) and contralateral tissue region from non-tumor hemisphere (lower panel). Green and magenta lines indicate the outer and inner diameter of the blood vessels respectively. Scale bar, 2µm. **D**, Measurement of the thickness of blood vessels in the tumor-bearing hemisphere of mice brains and contralateral region using Blood vessel ratio-SAE. Quantifications were performed in n=24 (WT), n=56 (KO), n=29 (normal brain WT) and n=44 (normal brain Esm1 KO). One way ANOVA followed by post hoc test, \*\**P=0.0030*. **E**, Proliferation of rEndocan treated (10ng/ml) and untreated TEC cells (TEC 15 and TEC15). Proliferation was measured on Day 5 post treatment, \*\**P=0.0012, ***P=0.0013*. **F**, Proliferation of TEC15 and TEC14 cells treated with conditioned media collected from the control GBM cells or GBM cells pretreated with rEndocan (10ng/ml) for 3 days. Proliferation was measured on day 5 post treatment. \**P=0.0173,* \*\*\**P<0.0002*. All quantitative data are average ±SD.

Consistent with these observations, qRT-PCR analysis demonstrated that vascular cell-associated genes known to play an integral part in maintaining barrier function, adhesion interactions, and tight junctions(*Podxl*, *Cld5*, and *Cdh5*)^24,25^, along with a marker for endothelial cells (*Pecam1*)^15^ were downregulated in tumors from *Esm1* KO mice (**Figure S2d**). Importantly, brain vascularization in the areas that did not contain tumor was not appreciably different between wild type and *Esm1* knockout mice, as can be seen from CD31 staining (**Figures s2c**) and electron microscopy (**Figures 2c-d**).

We then performed RNA-seq expression profiling of tumors formed by 7080 cells in WT and *Esm1* KO mice and identified 1449 downregulated and 283 upregulated genes (fold change >4; adjusted p-value < 0.05) in KO samples. Subsequent Gene Set Enrichment analysis (GSEA) revealed alterations in many important pathways including downregulation of the proliferation, DNA repair as well as Myc and VEGFA signaling in tumors formed in *Esm1* KO mice (**Figure S2e** and **Table S1)**.

Thus far, our findings indicate that Endocan promotes the hypervascularization phenotype of murine gliomas. This effect can be mediated either by autocrine (GBM cells promote Endocan expression in VE cells, which further enhance proliferation of VE cells) or paracrine (GBM cells treated with Endocan secrete other factors which promote vascular cell growth) mechanisms, or a combination of the two. In order to discriminate between these hypotheses, we first treated two human TEC cell lines with rEndocan and in both cases observed a significant increase in proliferation in response to Endocan (**Figure 2e)**. Next, to test the potential for an indirect effect of Endocan on endothelial cells, we treated GBM cells with rEndocan for 3 days, and then replaced the rEndocan containing media with regular GBM media. After 4 more days of incubation, conditioned media was collected and added to TECs. Results of this experiment demonstrated significantly higher proliferation of TECs in Endocan-induced conditioned media compared to the control media (**Figure 2f**).

Altogether our data suggest that Endocan is essential for the development of neovascularization in GBM. This protein, secreted by tumor-associated VE cells, can directly enhance proliferation of VE cells within GBM and also can stimulate GBM cells to secrete other factors additionally promoting vascularization of the malignancy.

### Endocan induces stable phenotypic alterations of GBM cells regulating their spatial identity

Next, we aimed to assess if GBM cells propagated *in vivo* in a microenvironment with or without Endocan acquire any stable differences in their phenotype. To answer this question, we generated sphere cultures from tumors formed in WT and *Esm1* KO mice brains (WTD and KOD cells respectively). Even under identical *in vitro* cultivation conditions, KOD GBM cells had reduced proliferation compared to WTD counterparts. Furthermore, KOD cells no longer responded to the addition of rEndocan (**Figure 3a**). To further characterize differences between these cells, we labeled WTD and KOD cells with GFP and mCherry, respectively, and co-injected them into the brains of *Esm1* WT mice (**Figure 3b**). IHC analysis of the resultant tumors demonstrated that WTD cells were located at the tumor edge region and infiltrated into the normal brain, whereas KOD cells appeared to be largely confined to the tumor core (**Figures 3c, d**). To quantitatively assess this phenomenon, we measured the number of cells of each type as a function of the distance from the injection site, demonstrating that KOD cells were mainly present in the tumor core-region, whereas WTD cells were found to be highly migratory and spread across the tumor and normal brain (**Figure 3e**). Interestingly, CD31 staining indicated that the core lesion predominantly formed by KOD cells was hypovascular, while the edge lesion created mainly by WTD cells was hypervascular (**Figure 3f**). The tumor core areas exhibited higher immunoreactivity to hypoxic and mesenchymal GBM markers (HIF1α, YKL-40, p-65 NF-κB and ALDH1A3)^26–29^ (**Figure 3f** and **Figure S3**), while the edge regions had higher levels of proneural markers PDGFRA and Olig2 (**Figure S3**). Taken together, these findings suggest that KOD cells have lost their capacity to promote neovascularization and to invade into the normal tissues even in the presence of Endocan.

**Figure 3:**
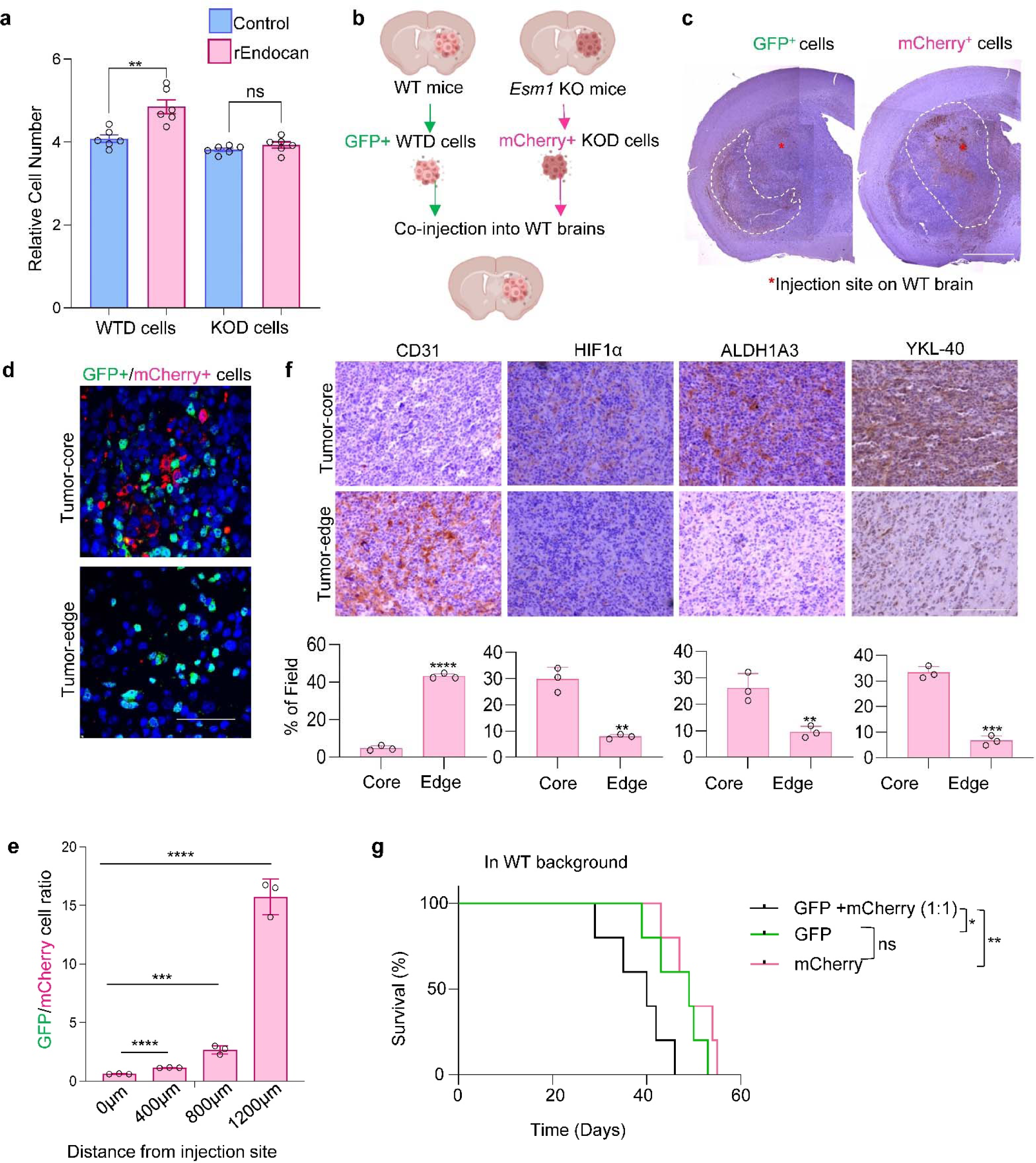
Endocan induces stable phenotypic alterations of GBM cells regulating their spatial identity. **A**, Proliferation assay of control and rEndocan treated (10ng/ml) WTD and KOD cells. *\*\*P=0.0023*, n=3 independent experiments. **B,** Schematic figure delineating the steps for the experiment. Briefly, mouse 7080 spheres were implanted into brains of WT and *Esm1* KO host, once tumors were formed, they were isolated into single cells, and were followed by labeling *Esm1* KO derived spheres with mCherry and WT derived spheres with GFP. Cells were then co-injected into WT host (n=5 mice). **C,** Representative IHC staining with anti-GFP (left) and anti-mCherry (right) antibodies of mice brains co-injected with WTD (GFP^+^) and KOD (mCherry^+^) cells (n=5 mice). The boundaries of the staining areas are shown by white lines, injection site indicated by red asterisk. Scale bar, 2mm. **D,** Representative IHF images of mice brain sections stained with anti-GFP (green), anti-mCherry (red) antibodies and DAPI (blue). Scale bar, 50 μm **E**, Ratio of GFP+ and mCherry+ cells in IHF images as in “D” at different distances from the site of injection. Quantification was performed using ImageJ software where mCherry+ cells and GFP+ cells were divided by DAPI staining. n=3 mice. *\*\*\*\*P<0.0001*; 800µm vs. 0µm, *\*\*\*P=0.0006*; 1200µm vs. 0µm, *\*\*\*\*P<0.0001*. **F,** Representative IHC images of core (upper panel) and edge regions (middle panel) of the tumor obtained as in “D” stained for CD31, HIF1α, ALDH1A3, and YKL-40. % of field positive areas for indicated proteins on lower panel (n=5 independent mouse samples). *\*\*P<0.01*, *\*\*\*P=0.0001*, *\*\*\*\*P<0. 0001.Scale* bar, 200µm. **G**, Kaplan-Meier survival analysis of *Esm1* WT mice intracranially injected with WTD and KOD cells or the mixture of both at 1:1 ratio. (n=5 mice per group), \**P*=0.047; ns. - non significant (*P*=0.31); \*\**P*=0.00642. *P*-values were determined by log-rank test. All quantitative data are average ±SD.

Surprisingly, we did not observe any survival differences between mice injected with WTD or KOD cells (**Figure 3g**). However, when a mixture of KOD and WTD cells was co-injected into the brain, animal survival was substantially reduced compared to the mice injected with either KOD or WTD cells alone (**Figure 3g**). We can speculate that the decreased survival of mice injected with the mixture of KOD and WTD cells may be explained by the increased intratumoral heterogeneity which by various mechanisms has been shown to promote GBM malignancy^28,30^.

Altogether, the striking differences observed between WTD and KOD cells suggestthat Endocan itself or the tumor microenvironment formed in the presence or absence of Endocan has a significant and long-lasting effect on the key vascular and infiltrative properties of GBM cells. Importantly, the phenotypical differences developed between cells propagated with or without Endocan can be stably maintained even if the cells are subsequently cultured under identical conditions *in vitro* or *in vivo*.

### Endocan binds to and activates PDGFRA in human glioblastoma cells

As a next step, we sought to understand the relevant mode of action of Endocan in glioblastoma. To identify possible receptors for Endocan on GBM cells, we purified integral membrane proteins from human gliomasphere line 157 and incubated these proteins with immobilized rEndocan. Subsequent elution of bound proteins and mass spectrometry identified PDGFRA as a potential Endocan-interacting protein (**Table S2**). To confirm this finding in a cell-free system, we tested the binding of the corresponding recombinant proteins. Pull down experiments demonstrated physical interaction between F_c_-tagged PDGFRA and His-tagged rEndocan (**Figure 4a)**. As another validation step, we used the proximal ligation assay^31^, which showed that rEndocan induces the dimerization of PDGFRA - the initial step in PDGFRA activation^32^ (**Figure S4a**). To further confirm the specificity and affinity of Endocan-interaction with PDGFRA, we performed a binding competition assay between rEndocan labeled with Alexa488 and recombinant PDGF-BB (one of the cognate ligands for PDGFRA), labeled with Alexa647. The addition of rPDGF-BB diminished rEndocan binding to glioblastoma cells in a concentration-dependent manner. Similarly, rEndocan decreased rPDGF-BB binding.

**Figure 4:**
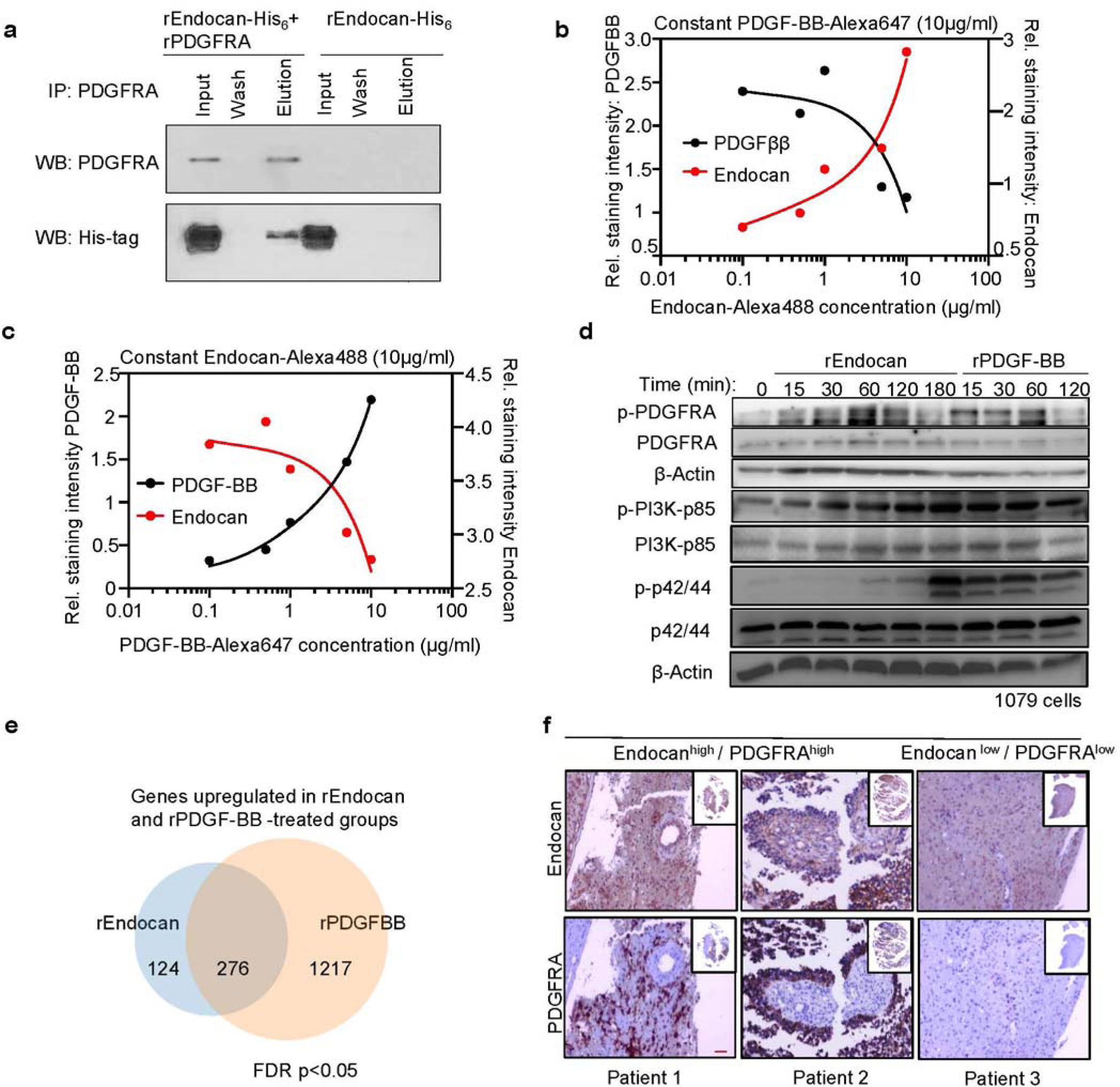
Endocan binds to and activates PDGFRA in human glioblastoma cells. **A,** Western blot (WB) analysis of samples obtained during pull down assay of His-tagged rEndocan and Fc-tagged rPDGFRΑ. **B, C** Competition binding assay of Endocan-Alexaflour488 and PDGF-BB-Alexaflour647. 1079 cells were incubated with indicated concentrations of both proteins and subsequently analyzed by FACS. **D**, WB analysis of 1079 cells incubated with 10ng/ml rEndocan or rPDGF-BB for the indicated period of time. **E,** Venn diagram showing genes which expression was altered by rEndocan, and rPDGF-BB after 3 days incubation as determined by RNAseq analysis in 413 cells. **F**, Representative immunohistochemical (IHC) images showing staining for Endocan (top panel) and PDGFRA (bottom panel) in three glioblastoma patients (Tissue Micro Array set generated from Osaka university cohort) Scale bar, 100µm. All quantitative data are average ±SD.

Importantly, a concentration of 10 ug/ml rEndocan was able to completely displace rPDGFββ from GBM cells, while rPDGF-BB at the corresponding concentration was able to displace only half of the rEndocan molecules (**Figures 4b-c**), suggesting that Endocan can interact with other receptors in addition to PDGFRA.

Given that PDGFRA activation by functional ligands induces receptor autophosphorylation, we investigated whether the addition of rEndocan alters the phosphorylation levels of PDGFRA in human gliomaspheres *in vitro* using PDGF-BB as a positive control^33^. We found that rEndocan (10 ng/ml, 400 pM) was able to induce appreciable autophosphorylation of PDGFRA in as early as 15 minutes and displayed similar temporal kinetics to that of rPDGF-BB (**Figures 4d** and **Figure S4b**). rEndocan treatment also resulted in the phosphorylation of the PDGFRA downstream targets such as p85α and PI3K^34^ along with phosphorylation of p44/42 MAPK^35^, further confirming the activation of the PDGFRA signaling pathway (**Figures 4d** and **Figure S4b**).

Next, we studied how Endocan and PDGF-BB affect GBM cells at the transcriptome level. RNA-seq analysis revealed that their effect on GBM cells was similar but not precisely the same (**Figure 4e**). More than 30% of Endocan-affected genes were not changed by the addition of PDGF-BB, indicating that while Endocan can activate the PDGFRA receptor, it might also have effects that are distinct from those of other PDGFR ligands, possibly due to the binding with other cellular receptors. Gene Set Enrichment Analysis (GSEA) revealed that Endocan specifically affects genes involved in angiogenesis, upregulation of Myc targets, and receptor tyrosine kinase (RTK) pathway activation (**Figure S4c**).

Finally, we evaluated the feasibility of Endocan and PDGFRA interactions in brain tumors using IHC. The staining of patient tumors demonstrated the presence of Endocan-expressing cells within perivascular areas that are adjacent to PDGFRA^+^ cells (**Figure 4f**).

Altogether, our data indicate that one of the molecular mechanisms by which Endocan affects GBM is binding to PDGFRA and subsequent activation of the downstream signaling pathways in tumor cells adjacent to VE cells in the microvasculature proliferation area.

### Endocan induces changes in the chromatin structure in the *Myc* promoter region

Since we demonstrated that GBM cells propagated in *Esm1* KO and WT mice showed substantial and highly persistent phenotypic differences, we hypothesized that Endocan-PDGFRA signaling can induce stable epigenetic alterations. To test this hypothesis, we performed ATAC-seq (Assay for Transposase-Accessible Chromatin sequencing) with *Esm1* WTD and KOD cells^36^. We identified 38 genes that had significantly different chromatin accessibility levels between samples, and among them, 9 genes showed different chromatin structure in their promoter regions (**Figures 5a and Table S3**). The strongest alterations were observed for the *Myc* promoter that showed more than 10-fold increased accessibility in WTD as opposed to KOD cells. To explore the relationship between open chromatin structure and gene expression levels, we next performed RNA-seq analysis of the same samples (**Figures 5b-c and Table S4**). Comparison of ATAC-seq and RNA-seq data revealed that among all genes with altered chromatin structure, *Myc* had the most substantial upregulation in the expression, with WTD tumors showing 25-folds higher level of *Myc* mRNA than KOD samples., IHC of WTD tumors revealed higher expression of c-Myc as compared to *Esm1* KOD samples (**Figure 5d**). To further confirm these data, we performed WB analysis for PDGFRA and Myc in the corresponding cells and saw consistent loss in the expression of both of these proteins in *Esm1* KOD samples (**Figure 5e**). To validate our finding that Myc is a downstream target of Endocan-driven PDGFRA signaling, we first treated our WTD and KOD cells with 10ng/ml of rEndocan for 3 days and found robust upregulation of *Myc* expression in WTD cells but not in *Esm*1 KOD cells (**Figure 5f**). Next, we used *in vitro* culture of patient-derived GBM cells (1079, 408 and 413) and confirmed that Myc was significantly upregulated at both the mRNA and protein levels in response to addition of the rEndocan to all tested cells (**Figures 5g-i, S5**).

**Figure 5:**
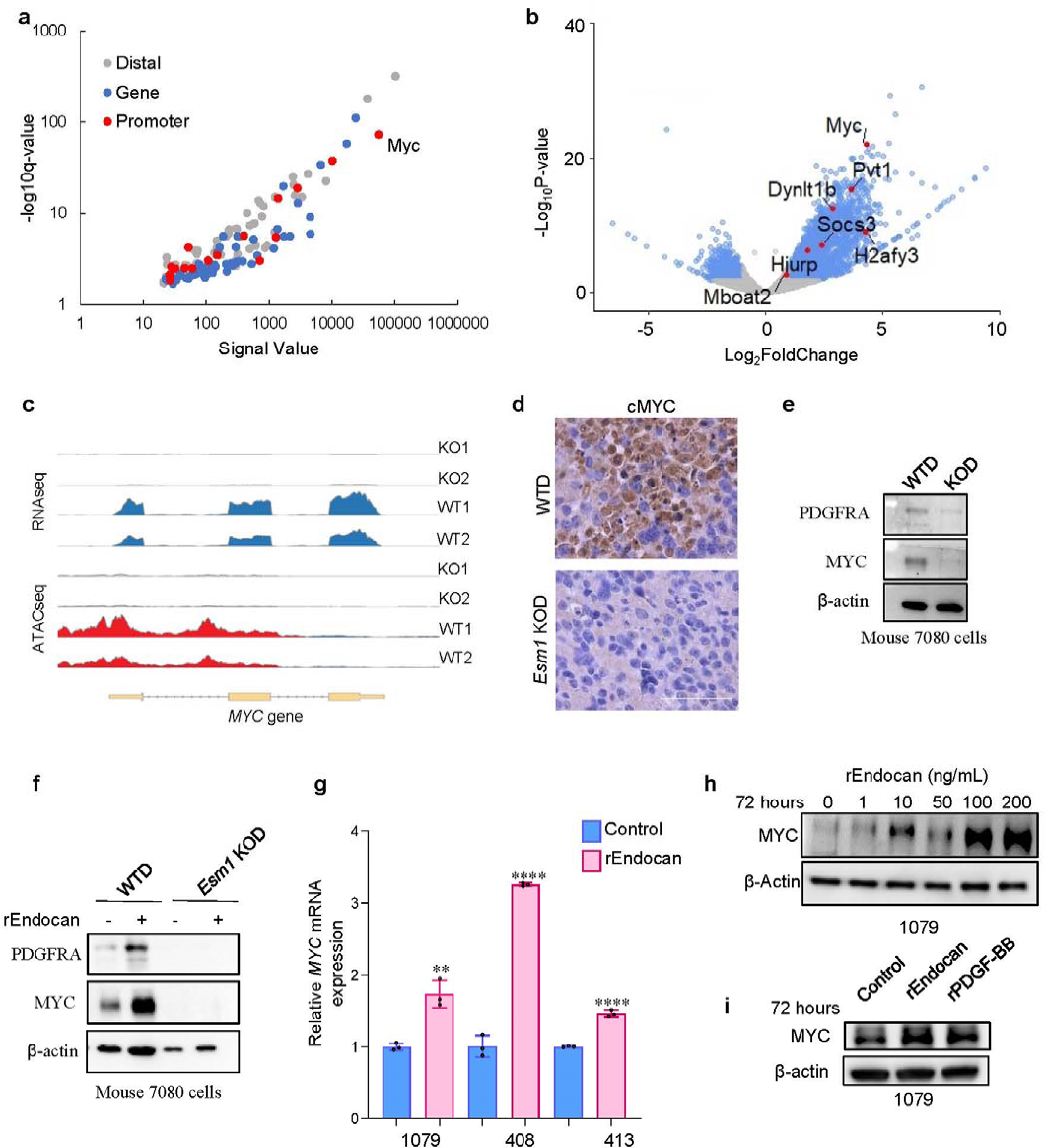
Endocan induces changes in the chromatin structure in the *Myc* promoter region. **A,** ATAC-seq analysis of tumors derived from *Esm1* WT and KO mice (n=2 mice per group). Individual dots represent peaks located in gene promoter region (red), in transcribed sequence (blue) and outside of the gene (gray). P-value of differences between *Esm1* WT and KO samples is indicated as well as normalized signal intensity. **B**, Volcano plot showing differentially expressed genes between tumors derived from *Esm1* WT and KO mice as determined by RNAseq analysis. (n=3 mice per group). Genes with altered chromatin structure of their promoter region as determined by ATAC-seq analysis are indicated with red dots. **C**, Sequencing read coverage of *Myc* gene as determined by RNA-seq (gene expression; blue) and ATAC-seq (chromatin accessibility; red) of the tumors as in “A” (n=2 mice per group). **D**, Representative microscopic images of WTD and KOD tumor slides stained with anti-MYC antibody. Scale bar, 100µm. **E**, Western blot analysis of WTD and KOD cells. **F**, Western blot analysis of WTD and KOD cells treated or untreated with rEndocan (10ng/ml) for 3 days. **G**, qRT-PCR analysis of *MYC* expression in control and rEndocan treated cells human GBM cells (1079, 408 and 413 cell lines). *\*\*P=0.003*, *\*\*\*\*P<0.0001*, n=3 independent experiments. **H**, Western blot analysis for MYC and β-actin protein in 1079 cells treated with increasing doses of rEndocan for 72 hours. **I**, Western blot analysis for MYC and β-actin protein in 1079 cells treated with rEndocan and rPDGF-BB for 72 hours. All quantitative data are average ±SD.

### Inhibition of the PDGFRA pathway attenuates the effect of Endocan on GBM cells

We next investigated whether the inhibition PDGFRA signaling would affect Myc expression. First, using lentiviral knockdown, we demonstrated that Myc level was substantially reduced upon depletion of PDGFRA (**Figure 6a**). Knockdown of PDGFRA also abrogated the proproliferative effect of Endocan on GBM cells (**Figure 6b**). Second, we utilized small molecule inhibitors ponatinib^37^ and nintedanib^38^ that have a high relative selectivity toward PDGFRA. Our results demonstrated that both compounds abrogate the effect of Endocan on PDGFRA activation and diminish Myc expression (**Figures 6c** and **Figure S6a, b**)., Treatment of GBM cells with ponatinib also reduced the proproliferative effect of rEndocan (**Figure 6d**).

**Figure 6:**
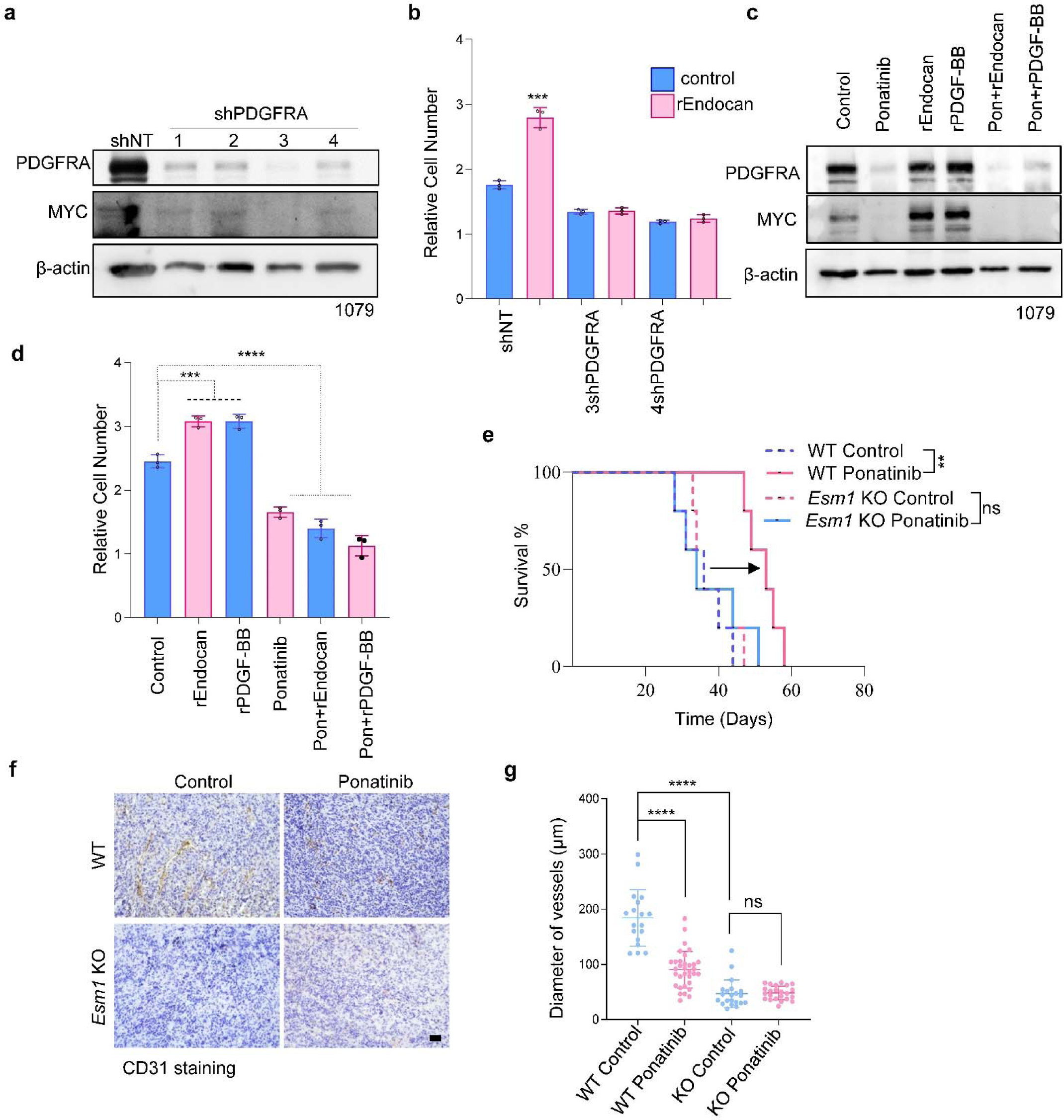
Inhibition of PDGFRA pathway attenuates effect of endocan on GBM cells. **A,** Western blot analysis of 1079 GBM cells after lentiviral mediated knockdown of PDGFRA with 4 different shRNAs against PDGFRA or Non Target shRNA (NT) as a control. **B**, Effect of rEndocan on proliferation of 1079 cells infected with shNT or shPDGFRA lentiviruses. n=3 independent experiments. *\*\*\*\*P<0.0001*. **C**, Western blot analysis of 1079 cells depicting PDGFRΑ, MYC, and β-actin protein level expression in cells treated with rEndocan, rPDGF-BB, ponatinib and their combinations for 72 hours. **D**, Effect of ponatinib on proliferation of 1079 cells incubated with rEndocan or rPDGF-BB. n=3 biological replicates, one-way ANOVA, post hoc T test. *\*\*\*P=0.0001*, *\*\*\*\*P<0.0001*. **E**, Kaplan Meier survival analysis of Bl/6 mice intracranially injected with WTD and KOD cells and treated with 50mg/kg/day dose of ponatinib for 5 days using oral-gavage. n=5 mice per group, *P*-value was determined by log-rank test; WT control vs. WT ponatinib, *\*\*P=0.00184*; KO control vs. KO ponatinib, *P=0.87, not significant*. **F**, Representative microscopic images of immunohistochemical (IHC) staining with anti-CD31 antibody of tumor sections obtained from mice as in “E”. **G**, Blood vessels density from tumor sections stained with CD31 antibody was analyzed using ImageJ software (n=20-33 vessels/group). Quantitative data are average ±SD; *\*\*\*\*P<0.0001*; *P=0.7852,* not significant. Scale bar, 100µm. All quantitative data are average ±SD.

Based on these data, we hypothesized that blockade of PDGFRA signaling in the WT background would attenuate Endocan-mediated activation of PDGFRA and subsequent Myc upregulation, leading to a hypovascular phenotype similar to that observed in *Esm1* KO tumors (**Figure 2**). To this end, we treated WT and *Esm1* KO mice bearing 7080 tumor cells with ponatinib. Consistent with our hypothesis, ponatinib enhanced survival only of the tumor-bearing WT, but not *Esm1* KO mice (**Figure 6e**), and reduced blood vessel formation, as can be seen from both CD31 immunostaining intensity (**Figure 6f**) and quantification of the blood vessels diameter (**Figure 6g).** Furthermore, IHC staining demonstrated that treatment with ponatinib resulted in a loss of Myc and Olig2 expression accompanied by increased level of p65-NFkB, a marker associated with tumor core (hypovascular) cells^26^ **(Figure S6c)**. Importantly, this effect was only observed in *Esm1* WT bot not KO mice.

Taken together, our findings suggest that Endocan-induced PDGFRA activation affects chromatin structure of the Myc promoter region leading to the prolonged upregulation of Myc expression and stable phenotypic alterations of GBM cells. Importantly, this pathway can be inhibited by small molecule compounds targeting PDGFRA. Tumors formed in the absence of Endocan utilize other mechanisms to drive their growth and are less sensitive to PDGFRA inhibitors.

### Endocan protects glioblastoma cells from radiotherapy

Previous studies have demonstrated that the tumor vascular niche provides protection to GBM cells against radiotherapy (IR)^39^. Therefore, we sought to determine the role of Endocan signaling in this process. First, we demonstrated that IR substantially upregulated *ESM1* transcription in VE cells *in vitro* as measured by qRT-PCR (**Figure S7a)** and increased secretion of Endocan protein by more than 3 folds as determined by ELISA (**Figure 7a**). Given these findings, we assessed whether IR-induced upregulation of Endocan could play a role in radioprotection of GBM cells by VE cells. First, we pretreated GBM cells with rEndocan or HBEC-5i conditioned medium (CM) for 3 days, followed by IR at 8 Gy. Analysis of gliomaspheres on day 5 post-irradiation revealed that both rEndocan and HBEC-5i CM significantly decreased IR-induced apoptosis (**Figure 7b** and **Figure S7b)**, prevented cell-cycle arrest (**Figure S7c**), and promoted GBM cell survival after radiation treatment (**Figures S7d)**. Furthermore, cells pretreated with Endocan had significantly reduced γ-H2AX staining intensity compared to the control groups (**Figure 7c**), indicating diminished IR-induced DNA damage^40^. Finally, given that radiation promotes mesenchymal differentiation of gliomaspheres^26^, we examined the effect of Endocan on the expression levels of mesenchymal (CD44) and proneuronal (CD133) GBM markers in irradiated 1051 cells. Pretreatment with either HBEC-5i CM or rEndocan suppressed radiation-induced downregulation of CD133 and upregulation of CD44 (**Figure S7e**), and therefore, protected GBM cells from mesenchymal differentiation.

**Figure 7:**
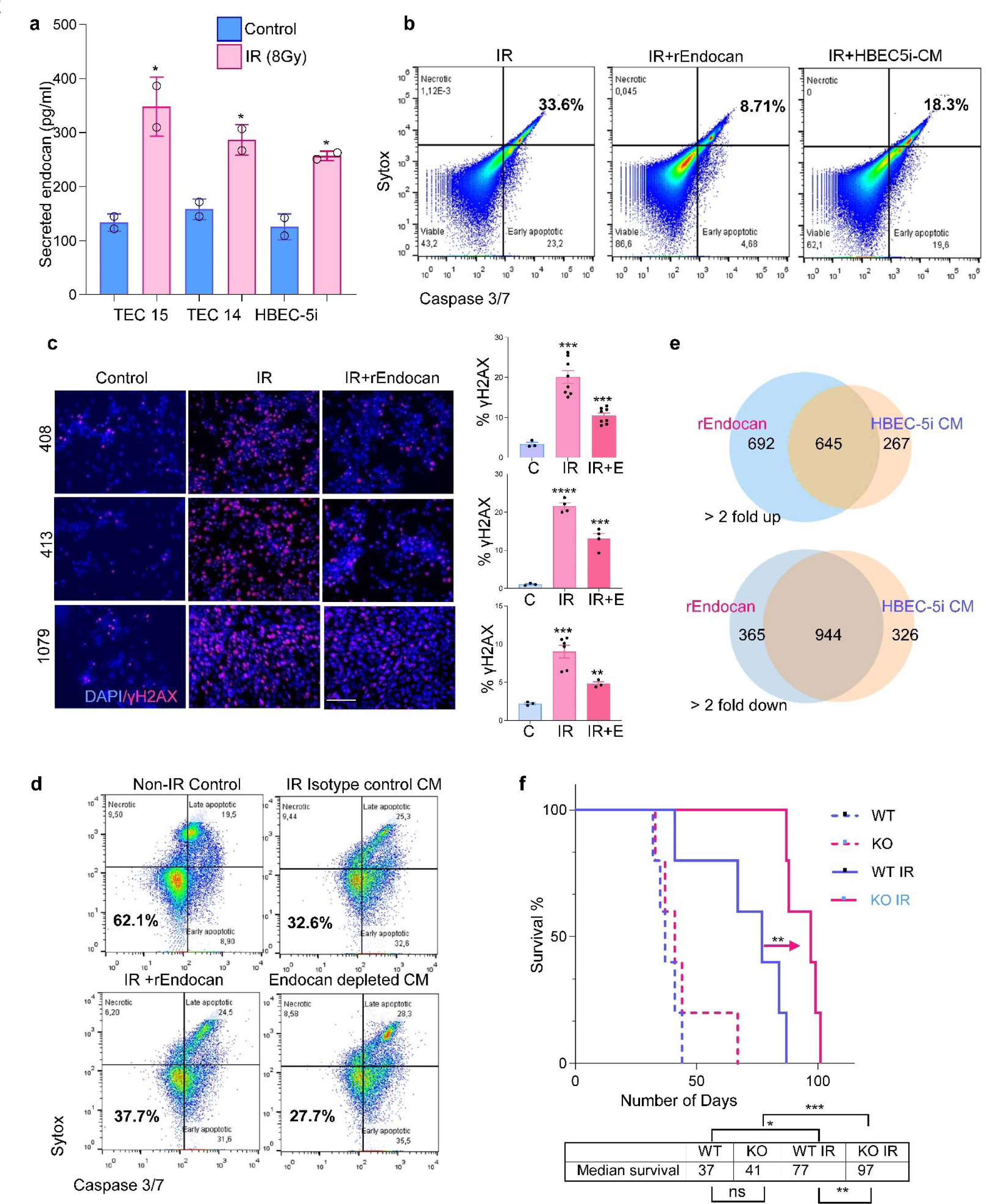
Endocan protects glioblastoma cells from radiotherapy. **A**, ELISA assay comparing levels of secreted Endocan in conditioned media collected from HBEC-5i cells, TEC14, and TEC15 cells following irradiation with 8 Gy at 48-hour post radiation time-point. n=2 independent experiments. *\*P=0.018, **P=0.0332*; *\*\*\*P=0. 0333*. **B,** Flow cytometry analysis of 711 cells treated with rEndocan or HBEC-5i CM and for 3 days and subsequently irradiated with 8 Gy. Samples were collected on Day 5 after irradiation and stained for caspase 3/7 and SYTOX. **C**, Representative microscopic images of GBM cells (408, 413, 1079) treated with rEndocan for 3 days and then treated with 8 Gy. Sham irradiated cells were used as control. Cells were fixed 16 hours after radiation and stained for γ-H2AX (Ser139) (left panel). Images were taken from multiple regions and n=3 independent replicates. % γH2AX positive cells were measured by dividing γH2AX cells by a total number of nuclei (DAPI positive cells) using ImageJ software (right panel). Scalebar: 125µm. *\*\*P=0.0012*, *\*\*\*P<0.001*, and *\*\*\*\*P<0.0001*. **D,** FACS analysis of Caspase 3/7 and SYTOX staining in 1051 GBM cell preincubated for 3 days with HBEC-5i CM that was incubated with Endocan blocking antibody (Endocan depleted CM), or isotype control antibody; cells preincubated with rEndocan was used as a positive control. After incubation cells were irradiated at 8 Gy and analyzed after 2 more days. **E**, Venn diagram representing differentially expressed genes in 1051 GBM cells irradiated with 8 Gy alone (control) or in a presence of rEndocan or CM from HBEC5i cells. Cells were collected 5 days after radiation. Genes which expression was altered at least 2 folds were used for the analysis. **F**, Kaplan Meier survival analysis of *Esm1* WT or *Esm1* KO mice intracranially injected with 7080 cells and subsequently irradiated with 3 doses of 2.5 Gy 14 days after injection. n=5 mice per group. *P*-value was determined by log-rank test, *ns – non significant*; *\*P=0.00875; **P=0.00443*; *\*\*\*P=0.00206*. All quantitative data are average ±SD.

To confirm that the protective effect of the secretomes from VE cells was specifically mediated by Endocan protein but not by other factors, we depleted Endocan from HBEC-5i CM using an anti-Endocan antibody. This depletion resulted in a decrease in the radioprotection of the HBEC-5i CM as measured by caspase 3/7 activity assay and a cellular proliferation assay (**Figure 7d**, **S7f**). RNA-seq of gliomaspheres that were pretreated with either HBEC-5i CM or rEndocan and subsequently exposed to 8 Gy IR demonstrated that rEndocan largely recapitulated the effects of HBEC-5i CM on the transcriptome of gliomaspheres after irradiation (**Figure 7e**). These findings suggest that Endocan is the major mediator for the VE cell-driven radioprotective effect on glioblastoma cells.

Finally, we investigated the effect of radiation treatment on survival of tumor-bearing *Esm1* WT and KO mice (**Figure 7f**). Both tumor models showed substantial shrinkage in size after radiation. However, radiation treatment resulted in a significantly greater enhancement of survival of tumor-bearing KO mice than WT mice (**Figure 7f, S7g**). These results are in a good agreement with our RNAseq data that demonstrated downregulation of DNA repair pathway in tumors formed in *Esm1* KO animal (**Figure S2e)**.

Altogether, our findings demonstrate that Endocan is a key mediator of the radioprotective effect of VE cells on GBM cells. Importantly, Endocan secretion is additionally increased upon IR which further promotes GBM cell viability and prevents radiation-induced mesenchymal differentiation of glioblastoma cells.

## Discussion

More than 50 years ago, it was demonstrated that malignant tumors promote microvascular proliferation, which in turn provides nutritional support that enables further cancer progression^41^. However, during the last decade, it has become clear that the relationship between tumor and VE cells is far more complicated. They form a sophisticated multilayered interaction network that shapes the key properties of both cell types. Although many components of this crosstalk have been identified, the key factors that orchestrate the entirety of tumor-vasculature signaling are not well elucidated. In the current study, we investigated some of the mechanisms of bidirectional interactions between GBM and VE cells. A dataset generated during our previous work allowed us to identify Endocan (*ESM1* gene) as the most significantly upregulated secreted protein in tumor-associated VE cells when compared to the normal brain endothelium. Our experiments revealed that on the one hand, Endocan affects the phenotype of GBM cells, and on the other hand, by direct and indirect mechanisms further promotes microvascular proliferation within the tumor (**Figure. 8**). Using *Esm1* KO mice, we demonstrated that the absence of Endocan has a dramatic effect on both GBM cells and tumor-associated vasculature, suggesting that this protein might be one of the master regulators of GBM-VE crosstalk.

**Figure 8:**
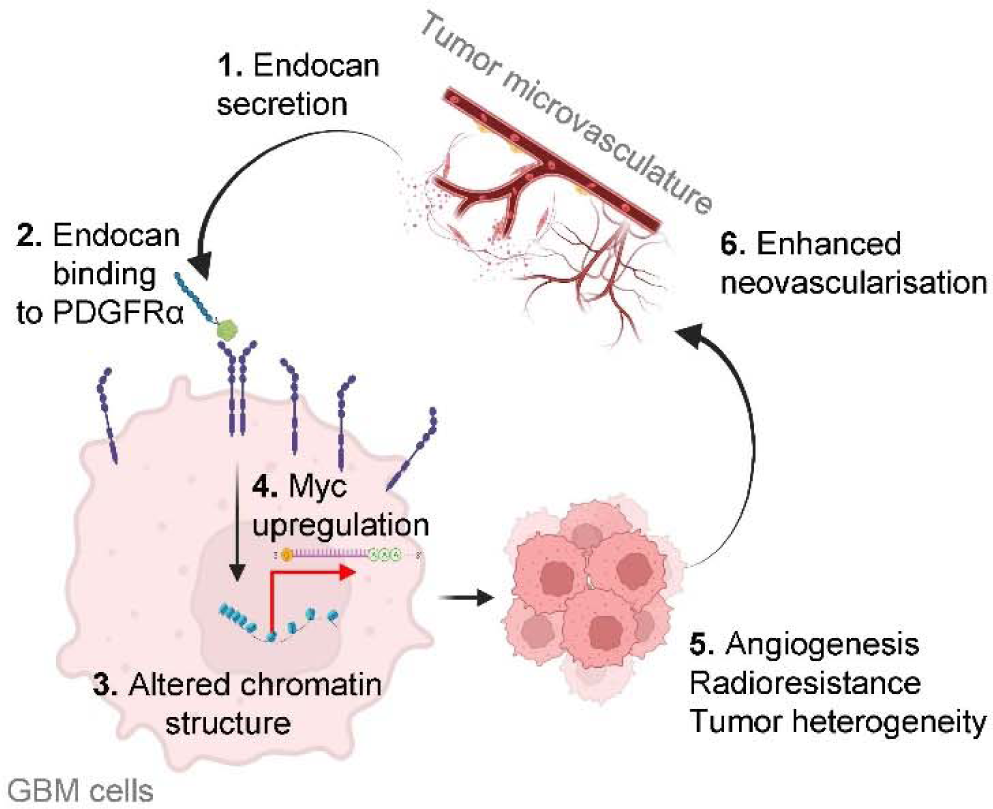
Schematic summary illustrating the role of Endocan in GBM-VE crosstalk. Endocan secreted by VE cells binds to PDGFRA receptor and activates downstream pathway, resulting in upregulated MYC expression and subsequent increase in GBM proliferation, radioresistance, intratumoral heterogeneity and further enhancement of angiogenesis.

A previous study demonstrated that Endocan and vascular endothelial growth factor A (VEGF-A) form a positive feedback loop in which VEGF-A promotes expression of Endocan in endothelial tip cells while Endocan further increases bioavailability of VEGF-A^13^. Considering that VEGF-A is one of the key vasculature growth factors secreted by cancer cells, Endocan upregulation in GBM-associated VE cells is not surprising. Interestingly, prior studies have also implicated Endocan in modulating cell proliferation, migration, and invasion in renal, colorectal, non-small cell lung, and hepatocellular cancers^10^, and showed that its expression correlated with poor prognosis in GBM patients^7^. Our data provide mechanistic explanations of these findings and suggest Endocan as a possible target for therapy as well as a potential diagnostic marker that may be used to predict GBM sensitivity to various types of treatment.

To determine the molecular mechanism by which Endocan affects GBM, we first identified its receptor on the surface of GBM cells. We demonstrated that Endocan binds to and activates PDGFRA which subsequently phosphorylates multiple downstream targets such as PI3K and ERK^42^. Endocan was previously shown to interact with CD11a integrin in human lymphocytes and Jurkat cells^43^ and that it could also activate EGFR signaling in non-small cell lung cancer^44^. However, our analysis did not identify either EGFR or CD11a as Endocan binding partners in GBM. It is possible that due to its unique structure that contains N-terminal 18 kDa protein component and C-terminal 30 kDa dermatan sulfate chain Endocan can interact with a high variety of cell surface receptors as well as extracellular matrix components^13^, and the net functional outcome of Endocan will depend on the expression level of its binding partners in addition to its relative affinity for them. This hypothesis is in good agreement with our data indicating that Endocan has a similar but not identical effect as PDGF-BB– one of the cognate ligands of PDGFRA, suggesting that Endocan may activate other yet unknown receptors in GBM. However, as we demonstrated that PDGFRA inhibition is sufficient to abrogate the effect of Endocan on GBM cells, we can speculate that, at least in our experimental model, PDGFRA is the main receptor for Endocan. Importantly, these data also provide insights into the non-canonical mechanisms of PDGFRA activation. Although the classical ligands for PDGFRs are members of PDGF family, other ligands for these receptors such as VEGF-A^45^ and CTGF^46^ have been identified. Therefore, Endocan may serve an additional PDGFR ligand allowing cancer cells to activate PDGFRA signaling pathway, even if PDGFs are absent or diminished Somewhat surprisingly, our results demonstrated that Endocan itself or microenvironmental conditions formed *in vivo* in the presence or absence of Endocan induce highly stable phenotypic alterations of GBM cells that persist even after months of cultivation in identical conditions. As this result was reproduced in two substantially different mouse GBM models, we hypothesized that it was associated with epigenetic alterations rather than with *de novo* mutations arising during propagation. Subsequent experiments revealed that while, overall, chromatin structure of GBM cells propagated in ESM1 WT and KO mice was very similar, there was a substantial difference in Myc promoter accessibilitydifference in the chromatin accessibility in the *Myc* gene.. Consistent with these findings, we demonstrated upregulation of Myc at both protein and mRNA levels. These results are in good agreement with previously published data indicating that PDGFR activation promotes Myc expression^47^ and that Myc level is regulated by the chromatin structure of its promoter region^48^. However, the exact molecular mechanism whereby PDGFR activation leads to altered chromatin and elevated Myc expression remains unknown.

As most of our experiments demonstrated that Endocan promotes aggressiveness of GBM cells, one could expect that GBM tumors developed in the presence of Endocan would show more malignant properties than the Endocan-deprived counterpart. Surprisingly, this was not the case. We demonstrated that GBM cells propagated in *Esm1* KO mice were as aggressive as the ones in *Esm1* WT. Nevertheless, our further experiments revealed that Endocan does increase the malignancy of GBM if cancer cells propagated in ESM1 WT and KO mice are co-injected together into the same animal. These findings are in good agreement with previous work indicating that a higher level of GBM intratumoral heterogeneity, rather than any specific tumor phenotype, correlates with worse patient outcome^49^. We can speculate that Endocan secretion restricted to the perivascular niche enhances intratumoral heterogeneity by promoting the development of two phenotypically distinct cell populations that grow in the presence or absence of Endocan. In our previous studies, we demonstrated that a variety of factors in the tumor necrotic core microenvironment promote mesenchymal-like transformation of GBM cells^28,50^. It is interesting to propose that the opposite process may exist in regions of microvascular proliferation, where Endocan protects GBM cells from mesenchymal differentiation and stimulates more proneural-like properties. This hypothesis is indirectly confirmed by the observation that Myc signaling is upregulated in proneural GBM tumors, while the mesenchymal ones exploit pathways related to inflammatory response and IL6/JAK/STAT3^51^. Therefore, we can argue that in different GBM regions various microenvironmental factors may differentially affect GBM phenotype, thereby promoting intratumoral heterogeneity and increasing GBM malignancy, and that Endocan protein plays an important role in this process.

We found that Endocan had a broad effect on GBM cell phenotype: it i) promoted cell migration; ii) enhanced proliferation; iii) increased resistance to therapy; iv) protected GBM cells from radiation induced mesenchymal differentiation; and v) stimulated neovascularization. Such multifunctionality can be explained by two scenarios. First, Endocan might interact with different receptors on GBM cells, activating various signaling pathways to produce the broad spectrum of observed effects. A second possibility is that Endocan affects a master regulator of GBM phenotype which is solely responsible for the observed downstream events. Discriminating between these possibilities will be important for the development of Endocan-targeted therapy, as in the first case it will be rational to inhibit Endocan protein itself, while in the second scenario, blocking of Endocan’s receptor or its downstream targets might be more beneficial.

While our current data do not allow us to definitively distinguish between these two mechanisms, the majority of our results suggest that most if not all effects of Endocan can be explained by the activation of the PDGFRA-Myc signaling axis. Indeed, Myc is known to act as a master regulator of angiogenic factors in GBM and other solid tumors^55^ and is also heavily involved in the regulation of GBM stemness, radioresistance and metabolisms^52,53^. However, further studies are needed to precisely dissect the molecular mechanisms underlying the effect of Endocan on GBM cells.

From the therapeutic perspective, blockade of Endocan signaling in GBM cells can be achieved by at least four different approaches: i) downregulation of Endocan secretion in VE cells; ii) removal of the already secreted Endocan protein from the extracellular space; iii) direct inhibition of PDGFRA; and iv) inhibition of Myc. Importantly, we demonstrated that treatment with a PDGFRA inhibitor, ponatinib, more substantially increased survival of animals from the *Esm1* wild-type group as opposed to the *Esm1-*KO mice, indicating that tumors that have a high level of expression of Endocan may be more sensitive to this type of treatment. On the other hand, we showed that the removal of Endocan from conditioned medium by antibodies increases the efficacy of radiotherapy *in vitro*. Consistent with these latter findings, a prior study demonstrated that treatment of mice with metastatic breast cancer tumors with 1-2B7, a high affinity anti-Esm1 monoclonal antibody, yielded an improved response to bevacizumab treatment i*n vivo*^54^. In an alternate approach, the addition of sunitinib (multi-tyrosine kinase inhibitor) to HUVEC endothelial cells has been recently shown to specifically prevent upregulation of Endocan, demonstrating another means to inhibit Endocan-tumor interactions. Finally, although no clinically successful small molecule Myc inhibitors have been developed thus far^55^ multiple approaches such as blocking peptides^56^ and exosome mediate knockdown^57^ have shown promising results in *in vivo* GBM models. It is important to note that global loss of *Esm1* does not cause a noticeable defect in the brain or other organs, so systematically diminishing or even eliminating tumor-associated Endocan could indeed be a promising strategy to control glioblastoma progression and recurrence.

Unfortunately, development of resistance to Endocan-targeting therapy can be expected. Thus, we demonstrated that in the absence of Endocan, GBM cells undergo reprogramming that can desensitize them to PDGFRA inhibition. This coincides well with a recently published study that showed that upon blockade of the PDGFR signaling pathway, GBM cells acquire stable and persistent phenotypic alterations allowing them to rely on PDGF-independent proliferation mechanisms^58^. Therefore, it is important to develop approaches targeting both Endocan-PDGFRA dependent and independent populations of GBM cells within the tumor.

In summary, the findings presented here provide insights into the mechanisms of intercellular crosstalk between GBM and VE. We found that the endothelial secreted Endocan protein has a strong effect on both VE and GBM cells and seems to play an important role in GBM progression. However, similar to the results earlier obtained in a clinical trial of the anti-angiogenic drug bevacizumab which failed to improve overall survival of GBM patients due to metabolic reprograming^59^, glioblastoma cells can easily adapt to the absence of Endocan by using alternative signaling pathways. Therefore, in order to achieve progress in the therapy of this devastating disease, it will be critical to reveal the molecular mechanisms underlying the high plasticity and adaptation capacity of GBM cells. Our results point to the alteration in chromatin structure of Myc as one of pathways possibly responsible for such adaptation.

## Materials and Methods

### Informed consent and ethics committee approvals

This study was conducted under protocols approved by the IRBs and IACUCs of University of California Los Angeles (UCLA), MD Anderson Cancer Center (MDA), and University of Alabama at Birmingham (UAB).

### Cell cultures

Human gliomasphere cultures were maintained in DMEM/F12 medium supplemented with 2% B27 supplement (% vol), 20 ng/ml bFGF, and 20 ng/ml EGF. The bFGF and EGF were added twice a week, and the culture medium was replaced every 7 days. Experiments with neurospheres/gliomaspheres were performed with lines that were cultured for fewer than 30 passages since their initial establishment. To obtain Human Tumor-Associated Endothelial cell culture glioblastoma tissues from patients were freshly isolated during surgery, manually dissociated into single cells, and subsequently sorted for CD31+ cells using magnetic beads (ThermoFisher Scientific). Following confirmation of CD31 expression, VE cells were grown in fibronectin-coated flasks with Endothelial cell growth media (ScienCell) containing 2% FBS, 1% Penicillin-Streptomycin solution and 1% Endothelial growth supplement (ScienCell). HBEC-5i (ATCC) were cultivated in DMEM/F12 medium containing 10% FBS, 1% Penicillin-Streptomycin solution and 1% Endothelial cell growth supplement. Expression of CD31 was periodically checked by flow cytometry. Spontaneous tumors formed in *PTEN*, *TP53*, *NF1* deleted mice^19^ were grown in neurosphere media as described previously. Freshly obtained mice tumor (PDGFB-overexpressing mouse GBM model) was kindly shared by Dolores Hambardzyuman lab (Emory University). Tumors were isolated to obtain single cells for injection into mice. HBEC-5i (ATCC) cells were cultivated in DMEM/F12 medium containing 10% FBS, 1% Penicillin-Streptomycin solution and 1% Endothelial cell growth supplement. STR analysis was performed to confirm cell identity (**Table S5**). The cell lines were tested negative for mycoplasma contamination.

### Genetically modified mice

*Esm1* KO (Bl/6) mice are a generous gift from Dr. Ralf H. Adams^13^. All animals were maintained in accordance with the National Institute of Health (NIH) Guide for the care and Use of Laboratory Animals and were handled according to protocols approved by the University of Alabama at Birmingham Institutional Animal Care and Use Committee. The genomic status of *Esm1* was confirmed by PCR according to the prior studies.

### *In Vivo* Intracranial Xenograft Tumor Models

6-8-week-old NOD SCID mice (Prkdc^scid^) were used for intracranial tumor formation. In each mouse, 5 × 10^5^ cells of human patient-derived glioblastoma cells were injected Detailed protocol is described previously^7,14^.

### *In Vivo* Syngeneic Intracranial Tumor Models

Our general protocol for the intracranial tumor models was described previously^14^. Briefly, 5 × 10^5^ murine glioblastoma cells were injected into the brains of Bl/6 or *Esm1* KO mice. Mice were monitored and sacrificed when neuropathological symptoms developed. For immunohistochemical studies, mice were perfused with ice-cold PBS, followed by 4% paraformaldehyde (PFA). Mice brains were dissected and fixed in 4 % PFA solution for 48 hours and then transferred to 10% formalin for 48 hours. For tumor collection to perform RNA extraction, mice were sacrificed, and tumors were isolated and fast-frozen using liquid nitrogen.

### In vitro and in vivo irradiation

Cells or mice were irradiated at room temperature using X-ray irradiator (Gulmay Medical Inc., Atlanta, GA) at a dose rate of 5.519 Gy/min for the time required to apply an 8Gy dose. The X-ray beam was calibrated using NIST-traceable dosimetry and operated at 300kV and hardened using a 4mm Be, a 3mm AI, and a 1.5mm Cu filter. Mice were anesthetized prior to the irradiation. The body was covered with lead shield to avoid whole body irradiation.

### *In vivo* drug treatments

Mice were injected with 7080 cells to form intracranial tumors. On day 14 post injection, mice were given daily administration of 50mg/kg/day of Ponatinib (Selleckchem) by oral gavage for 5 days.

### Endocan enzyme-linked immunosorbent assay (ELISA)

The concentration of secreted Endocan was measured in conditioned medium using the Endocan ELISA kit (Boster Bio) according to the manufacturer’s protocol.

### Protein labeling

Recombinant Endocan and PDGF-BB were labeled with Alexa Fluor 488 Microscale Protein Labeling Kit (Thermo Fisher Scientific) and Alexa Fluor™ 647 Microscale Protein Labeling Kit (Thermo Fisher Scientific) respectively according to the manufacturer’s protocol. Labeled proteins were added to glioblastoma cells in different concentrations and after 30 min incubation on ice, cells were washed twice with PBS and analyzed by Attune NxT Flow Cytometer (Thermo Fisher Scientific). The obtained data were processed with FlowJo 10 software.

### Recombinant protein pull-down assay

To obtain a protein complex, recombinant His-tagged Endocan (R&D) was immobilized on 50 μl of HisPur Ni-NTA Magnetic Beads (Thermo Fisher Scientific) according to the manufacturer’s protocol. The beads were washed 3 times with PBS and incubated for 2 hours with Fc-tagged recombinant PDGFRΑ (R&D) under constant agitation. Next, beads were washed 3 times with PBS and bounded proteins were eluted with 300 mM imidazole in PBS and subjected to subsequent western blot analysis.

### Proximal Ligation Assay

Glioblastoma cells were plated in wells of Lab-Tek II chamber pre-coated with laminin and treated with Endocan or PDGFBB for 90 minutes. Next, cells were washed 3 times with phosphate-buffered saline (PBS) and fixed with 4% PFA in PBS for 15 min at room temperature. Cells were washed 2 times with PBS and permeabilized with 0.2% Triton-X100 in PBS for 15 minutes. All subsequent procedures were performed using “Duolink In Situ Orange Starter Kit” (Duolink) according to the manufacturer’s protocol.

### Mass spectrometry

Recombinant His-tagged Endocan (R&D) was immobilized on 50 μl of HisPur Ni-NTA Magnetic Beads (Thermo Fisher Scientific) according to the manufacturer’s protocol. The beads were washed 3 times with PBS and incubated with the fraction of plasma membrane proteins that were isolated from g1079 cells as described previously^60^ and solubilized in lysis buffer (50 mM Tris-HCl, 150 mM NaCl, 1% Triton X100, 0.1% sodium deoxycholate, protease inhibitor cocktail, pH 7.5). Next, beads were washed once with lysis buffer and 3 times with PBS and bounded proteins were eluted with buffer containing 8M Urea, 2M Thiourea, 10 mM Tris (pH=8). Protein concentrations were determined using the QuickStart Bradford protein assay (Bio-Rad) according to the manufacturer’s protocol.Eluates from immunoprecipitations were subsequently incubated with 5mM DTT at RT for 40 min. Then proteins were alkylated with 10 mM Iodoacetamide at RT for 20 min in the dark. Alkylated samples were diluted by the addition of 50 mM Ammonium Bicarbonate solution at a ratio of 1:4. Next, trypsin (0.01 μg per 1 μg of protein) was added, and the samples were incubated at 37°C for 14 h. After 14 h the reaction was stopped by the addition of Formic acid to the final concentration of 5%. The tryptic peptides were desalted using SDB-RPS membrane (Sigma), vacuum-dried, and stored at −80°C before LC-MS/MS analysis. Prior to LC-MS/MS analysis samples were redissolved in 5% ACN with 0.1% TFA solution and sonicated. Proteomic analysis was performed using a Q Exactive HF mass-spectrometer. Samples were loaded onto 50-cm columns packed in-house with C18 3 μM Acclaim PepMap 100 (ThermoFisher), with an Ultimate 3000 Nano LC System (ThermoFisher) coupled to the MS (Q Exactive HF, ThermoFisher). Peptides were loaded onto the column thermostatically controlled at 40°C in buffer A (0.2% Formic acid) and eluted with a linear (120 min) gradient of 4 to 55% buffer B (0.1% Formic acid, 80% Acetonitrile) in A at a flow rate of 350 nl/min. Mass spectrometric data were stored during automatic switching between MS1 scans and up to 15 MS/MS scans (topN method). The target value for MS1 scanning was set to 3·10^6^ in the range 300–1200 m/z with a maximum ion injection time of 60 ms and a resolution of 60000. The precursor ions were isolated at a window width of 1.4 m/z and a fixed first mass of 100,0 m/z. Precursor ions were fragmented by high-energy dissociation in a C-trap with a normalized collision energy of 28 eV. MS/MS scans were saved with a resolution of 15000 at 400 m/z and at a value of 1·10^5^ for target ions in the range of 200-2000 m/z with a maximum ion injection time of 30 ms.

Raw LC-MS/MS data from Q Exactive HF mass-spectrometer were converted to. mgf peaklists with MSConvert (version 3). For this procedure we use the following parameters: “--mgf --filter peakPicking true [1,2]”. For thorough protein identification in samples from immunoprecipitations, the generated peak lists were searched with MASCOT (version 2.5.1) and X! Tandem (ALANINE, 2017.02.01) search engines against UniProt human protein knowledgebase with the concatenated reverse decoy dataset. The precursor and fragment mass tolerance were set at 20 ppm and 0.04 Da, respectively. Database-searching parameters included the following: tryptic digestion with one possible missed cleavage, static modification for carbamidomethyl (C), and dynamic/flexible modifications for oxidation (M). For X! Tandem we also selected parameters that allowed a quick check for protein N-terminal residue acetylation, peptide N-terminal glutamine ammonia loss, or peptide N-terminal glutamic acid water loss. Result files were submitted to Scaffold 4 software (version 4.0.7) for validation and meta-analysis. We used the local false discovery rate scoring algorithm with standard experiment-wide protein grouping. For the evaluation of peptide and protein hits, a false discovery rate of 5% was selected for both. False positive identifications were based on reverse database analysis. We also set protein annotation preferences in Scaffold to highlight Swiss-Prot accessions among others in protein groups.

### *In vitro* migration experiment

Patient-derived Glioma cells (1079, 413, 408) were used for encapsulation. Approximately 600k cells per well were seeded into Aggrewell well plates (Stemcell Technologies) one day prior to encapsulation. Next day, the spheroids were encapsulated in hydrogels using a previously described hydrogel fabrication protocol^7^. Cell migration was observed at Day 1, 3, 6 and 9 post encapsulations by acquiring phase contrast images on a Zeiss Axio.Z1 Observer microscope with a Hamamatsu Orca Flash 4.0 V2 Digital CMOS Camera and Zeiss ZEN 2 (Blue Edition) software.

### Transmission Electron microscopy

Tissue from tumor-bearing mice were resected after anesthetizing and perfused with 1X PBS. Tissues were fixed in 4% paraformaldehyde and 2% glutaraldehyde. Next, the tissues were fixed with 1% Osmium tetroxide followed by dehydration in different concentrations of ethanol, and resin embedding. Following the embedding, sections were cut to 60-90 nm thickness, and then were counterstained with Uranyl Acetate and Lead citrate for imaging and analysis. Hitachi® H-7500 electron microscope was used for imaging and images were captured with NanoSprint12 AMT® camera. The outer-inner blood vessel ratio was calculated based on the inner-to-outer diameter of the blood vessel wall. Previously published Shape adjusted ellipse (SAE) method was used to calculate the diameters^23^. The SAE approach corrects for dispersion angles. One way ANOVA followed by post hoc test was performed, \*\**P=0.0030* (tumor bearing WT vs. *Esm1* KO) groups.

### ATAC sequencing and analysis

WTD and KOD tumor cells were processed for ATAC sequencing and repeated at least 2 independent times as previously described^39^. Briefly, 50,000 cells were washed with 50mL ice cold PBS and re-suspended in 50mL lysis buffer (10 mM Tris-HCl pH 7.4, 10 mM NaCl, 3 mM MgCl_2_, 0.2% (v/v) IGEPAL CA-630). The suspension was centrifuged at 500g for 10 minutes at 4 degrees. Samples were added with 50mL trans-position reaction mix of Nextera DNA library preparation kit (FC-121-1031, Illumina). DNA was amplified by PCR and incubated at 37°C for 30 minutes. MinElute Lit (Qiagen) was used to isolate DNA. NextSeq 500 High Output Kit v2 (150 cycle, FC-404-2002, Illumina) was used to sequence ATAC library. For analysis, alignment was carried out using the Burrows-Wheeler Aligner mem, peak calling using MACS2 (with parameter setting –nomodel –shift 75), GO enrichment analysis and peak annotation using HOMER and Enrich R (https://maayanlab.cloud/Enrichr/). UCSC Genome Browser was used to determine whether open regions displayed H3K27Ac and conserved TF binding sites. Integrated Genome viewer (IGV) was utilized to represent the open/peak regions in the experiments.

### RNA sequencing

cDNA were used in the library preparation using Ovation® Ultralow Library Systems (NuGEN) and samples were sequenced using an Illumina HiSeq 2000 sequencer (Illumina, San Diego, CA) in high output mode across 9 lanes of 50bppaired-end sequencing, corresponding to 4.3 samples per lane and yielding ∼45 million reads per sample. Additional QC was performed after the alignmentTotal counts of read-fragments that were aligned to all the candidate gene regions were derived using HTSeq program (www.huber.embl.de/users/anders/HTSeq/doc/overview.html) with Human Hg38 (Dec.2014) RefSeq (refFlat table) as a reference and used as a basis for the quantification of gene expression. Only uniquely mapped reads were used for subsequent analyses. Differential expression analysis was conducted with R-project and the Bioconductor package edgeR. Statistical significance of the differential expression, expressed as Log_2_ Fold Change (logFC), was determined, using tag-wise dispersion estimation, at p-Value of <0.005 unless stated otherwise. FPKM values were reported as a measure of relative expression units.

### Flow cytometry

For CD44 staining gliomaspheres were dissociated into single cells and stained with anti-CD44-APC antibody (Miltenyi Biotec) according to manufacturer’s protocol. For CD133 staining, gliomaspheres were dissociated into single cells and stained with anti-CD133-FITC (Biolegend) according to manufacturer’s protocol. For apoptosis assay, cells were stained with CellEvent Caspase-3/7 Green Flow Cytometry Assay Kit (ThermoFisher Scientific) according to the manufacturer’s protocol. To estimate the percentages of cells in the different phases of the cell cycle, propidium iodide (PI) dye (ThermoFisher Scientific) was used. Cells were fixed in 70% ethanol and treated with riboneuclease as recommended by the manufacturer’s protocol. All samples were analyzed by Attune NxT Flow Cytometer (Thermofisher Scientific) and the data were processed with FlowJo 10 software.

### Endocan blocking antibody experiment in HBEC-5i CM

For Endocan blocking experiments, HBEC-5i cells were grown without serum and 10ul of anti-Endocan antibody (ab103590) was added to 10ml of CM. 10 ug of control rabbit IgG was added to another 10 ml of CM. CM was incubated 2 hours at room temp with slow agitation followed by incubation with 100ul of ProtA/G magnetic beads (Thermofisher Scientific) for 1 hour. CM was then spun at 100g for 5 minutes and supernatant was carefully removed without disturbing the beads and subsequently filtered through 0.22µm filter.

### Cell proliferation assay

AlamarBlue reagent (Thermo Scientific) or Cell-Titer Glo (Promega) was used to determine the cell number under various treatments. Briefly, cells were seeded at a density of 5,000 cells per well in 96-well plates. AlamarBlue reagent was added into each well and fluorescence was measured (Excitation 515-565 nm, Emission 570-610 nm) using Synergy HTX multi-mode reader (BioTek). CellTiter-glo was used at 1:1 ratio and measured using a luminometer plate reader. Readings were taken on Day 0 and Day 3 or Day 5 after treatment.

### Western blot

The cell lysates were prepared in RIPA buffer containing 1% protease and 1% phosphatase inhibitor cocktail on ice. The sample protein concentrations were determined using the Bradford method. Equal amounts of protein lysates (10 μg/lane) were fractionated on a NuPAGE Novex 4%–12% Bis-Tris Protein Gel (Thermo Fisher Scientific) and transferred onto a PVDF membrane (Thermo Fisher Scientific). Subsequently, the membrane was blocked with 5% skim milk or 5% BSA for 1 hour and then incubated with the appropriate antibody at 4°C overnight. Membranes were then washed with 3X with TBST buffer and incubated with appropriate HRP-conjugated secondary antibodies (GE Healthcare) for 1 hour at RT. Protein expression was visualized with an Amersham ECL Western Blot System (GE Healthcare Life Sciences). β-Actin served as a loading control. ImageJ software (NIH) was used to analyze the Western blot results.

### Immunohistofluorescence

Immunohistofluorescence (IHF) staining was performed as described previously^14^. Briefly, tumors embedded in paraffin blocks were deparaffinized using Xylene and hydrated through 100%, 95% and 75% ethanol gradient. Antigen was retrieved using DakoCytomation target retrieval solution pH 6 (Dako). Samples were then blocked with serum-free protein block solution (Dako) and incubated with corresponding primary antibodies at 4°C overnight. Next, slides were incubated with Alexa Flour-conjugated secondary antibody for 1 hr at room temperature and mounted in Vectashield mounting medium containing DAPI (Vector Laboratories). Nikon A1 Confocal microscope (Nikon) was used to capture images.

### γ-H2AX assay

Chamber slides or wells were covered with laminin for 24 hours prior to cell seeding. Cells were allowed to grow and acclimate with treatment conditions. At the indicated times, the cells were irradiated or treated. Cells were then fixed in 4% paraformaldehyde, permeabilized with 0.5% Triton-X/PBS, and stained with an anti γ-H2AX antibody (1:500) and anti-rabbit conjugated Alexa 594 secondary antibody was used along with DAPI. Plates were imaged using EVOS microscope and ImageJ was used to manually quantify positively stained cells. At least 10-15 images were taken at 20X and in different areas per group.

### Immunohistochemistry

Immunohistochemistry (IHC) staining was performed as described previously^28^. Briefly, tumors embedded in paraffin blocks were deparaffinized using Xylene and hydrated through 100%, 95%, and 75% gradient of ethanol. Slides were then microwaved in the presence of DakoCytomation target retrieval solution pH 6 (Dako). Slides were incubated with 0.3% hydrogen peroxide solution in methanol for 15 minutes at room temperature to inhibit internal peroxide activity. Slides were then blocked with serum-free block solution (Dako) and incubated with corresponding primary antibody overnight at 4°C. Next, samples were incubated with EnVision+ System-HRP labeled Polymer (Dako) and visualized with DAB peroxidase substrate kit (Vector Laboratories). Images were captured using EVOS® FL inverted microscope (Thermo Fisher Scientific IHC scoring was performed using a previously described method. Vessel quantification for IHC images were measured using Vessel Quantify_IHC and IHC profiler Plugin in ImageJ, for CD31 staining in tissues^7,61^).

### Lentivirus production and transduction

Lentiviruses were produced as described before^28^. Briefly, HEK293FT packaging cells (Invitrogen) were co-transfected with lentiviral vectors encoding shRNAs or GFP or mCherry and two packaging plasmids psPAX2 and pMD2.G. Growth media was changed the following day and lentiviruses-containing supernatants were harvested at 72 hr after transfection and concentrated 100-fold using Lenti-X concentrator (Clontech). For infection, GBM cells were dissociated into single cells with accutase and seeded on laminin coated 6 well plates at 8×10^5^ cells per well. Next day, 8µg/ml of Polybrene (EMD Millipore) along with viral supernanatant were added to GBM cells. Two days after infection, transduced cells were selected with 1µg/ml of puromycin (Sigma) for 3 days.

### Tissue Microarray

Tissue microarray consisting of 0.6-mm cores from formalin-fixed, paraffin-embedded tissue blocks were generated using patient derived glioblastoma tissue samples at the Osaka City University (Cohort #3 n=38) after approval of the experimental protocol by the Institutional Review Board.

### RNA Isolation and Quantitative Real-time PCR

mRNA was extracted and purified using the RNeasy Mini kit (Qiagen) according to the manufacturer’s protocol. Nanodrop 2000 spectrophotometer was used to determine the concentration and quality of RNA. RNA (0.5-1µg) was reverse-transcribed in cDNA using iScript Reverse Transcription Supermix (Bio-Rad) and then amplified using the following cycling conditions; 95°C for 5 min, and then 50 cycles of 95°C for 20 s, 60°C for 20 s and 72°C for 20 s. qRT-PCR was performed on StepOnePlus thermal cycler (Thermo Scientific) with SYBR Select Master Mix (Thermo Scientific). 18s, B-actin was used as an internal control.

### Quantification and Statistical analysis

GraphPad Prism 10 was used for statistical tests and all details can be found in figure legends. Reported experiments were conducted at least three independent times unless stated otherwise. Error bars on all the graphs show Mean and Standard deviation (SD). Statistical analysis was performed using using the unpaired student’s t-test unless mentioned otherwise in figure legends. One-way ANOVA followed by post-hoc t test was used for statistical comparisons of multiple groups. Log-rank analysis was used to determine the statistical significance of Kaplan-Meier survival curves. *P* values less than 0.05 were considered to be significant. No samples, mice or data points were excluded from the analysis reported.

## Data availability

All raw RNA-seq data will be available in the NCBI Gene Expression Omnibus (prepared prior to publication).

## Acknowledgements

We would like to express our sincere appreciation to all the patients and families, who kindly allowed us to obtain their tumor samples for this study. We thank Dr. Ralf Adams (Max Planck Institute for Molecular Biomedicine) for generously sharing cryopreserved *Esm1* embryos. Lastly, we acknowledge the contribution by all the members in the Nakano and Kornblum laboratories (past and present) for technical assistance.

## Author Contribution

Conceptualization: S.B, M.S.P, IN, H.I.K

Methodology: S.B, M.S.P, N.P.B, L.A.H, A.B.H, S.K.S, S.A.G, I.N, H.I.K

Formal Analysis: S.B, M.S.P, I.N, H.I.K

Laboratory investigation: S.B, M.S.P, N.S, M.S.K, N.P.B, L.A.H, A.S, S.Y, Y.G, S.D.M, D.Y, J.A.O, B.T, A.L.B

Bioinformatics analysis: K.S.A, R.K, Y.Q Writing – Original Draft: S.B, M.S.P, I.N, H.K

Writing – Review & Editing, S.B, M.S.P, M.A.N, N.P.B, L.A.H, I.N, H.I.K

Funding Acquisition: M.S.P, K.S.A, A.B.H, L.A.H, A.L.B, S.A.G, I.N, H.I.K

Resources: H.I.K, A.L.B, I.N

Supervision: H.I.K, I.N

---

**Supplementary Figure 1:**
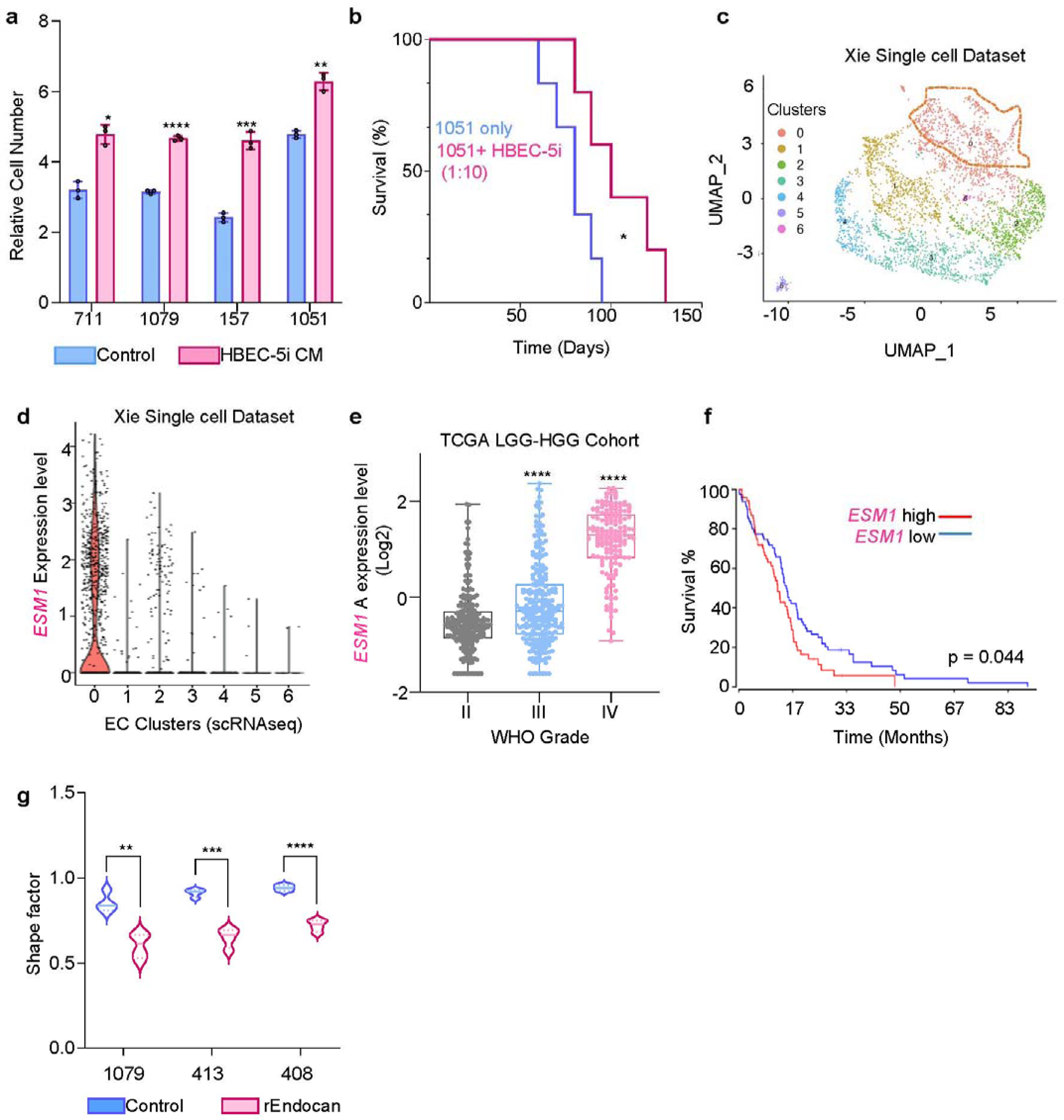
**A,** Proliferation of GBM cells (711, 1079, 157, 1051) treated with control media or conditioned media collected from Human Brain Endothelial Cells-5i (HBEC-5i) cells; n=3 independent experiments. \**P=0.0017*, \*\**P=0.0007,* \*\*\**P=0.0002*, and \*\*\*\**P<0.0001*. **B,** Kaplan-Meier survival analysis of SCID mice intracranially injected with gliomaspheres alone (1051) or gliomaspheres with VE cells (1051+HBEC-5i) at a 10:1 ratio. (n=5 mice per group). Log-rank test, \**P=0.0323*. **C,** Analysis of Xie single cell RNAseq dataset^15^ showing different clusters of VE cells present within the GBM tumor. Cells related to cluster 0 are highlighted. **D**, Analysis of *ESM1* expression in clusters shown in “C”. **E**, Expression of *ESM1* in TCGA’s WHO grade II, III, and IV glioma tumors. One-way ANOVA followed by Tukey’s post hoc test; *\*\*\*\*P<0.0001*. **F**, Kaplan-Meier analysis showing the survival of 667 glioma patients subdivided based on *ESM1* expression level. Log-rank test; \**P = 0.044*. **G**, Quantification of shape factor/circularity in rEndocan treated and control group of spheroids formed by 408, 413 or 1079 cells. Circularity was calculated using ImageJ shape description. n=24 spheroids for independent experiments, \*\**P*=0.0016; \*\*\**P*=0.0001; \*\*\*\**P*<0.0001. All quantitative data are average ±SD.

**Supplementary Figure 2:**
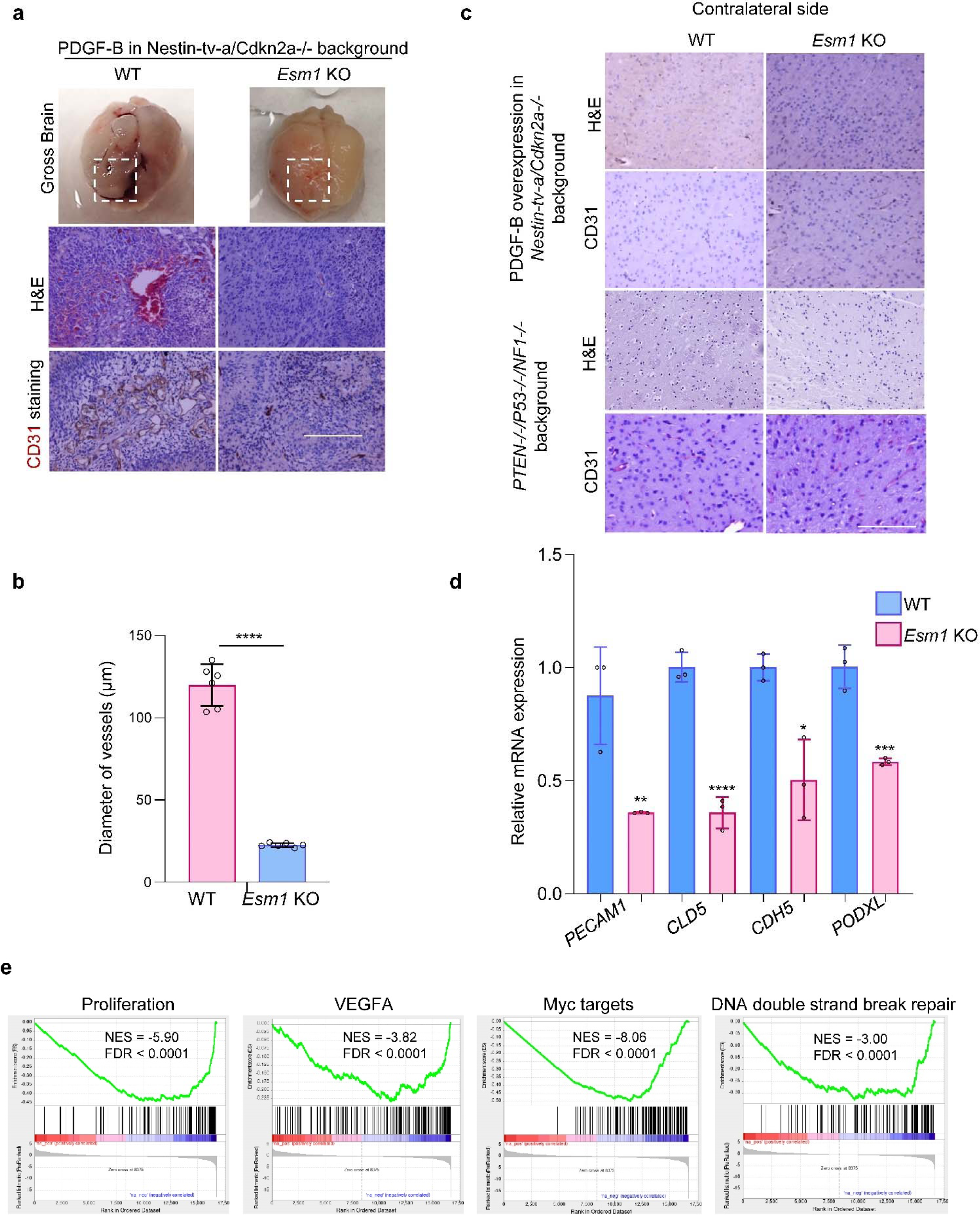
**A,** Tumors formed by murine glioblastoma cells (isolated from Nestin-Tva/Cdkn2a-/- mice intracranially injected with RCAS-PDGFB lentiviruses) in WT and *Esm*1 KO mice brains. Images of the brains (upper panel); H&E staining of the tumor slices (middle panel, Scale bar, 200 μm); CD31 staining (lower panel, Scale bar, 200 μm). **B**, Quantitation of the diameter of the blood vessels in samples as in “A”. n=5 mice brains, *\*\*\*\*P<0.0001*. **C,** H&E staining (upper panel) and CD31 staining (lower panel) of non-tumor/contralateral brain sections obtained from tumor bearing *Esm1* WT and KO mice injected with cells related to two different mouse GBM models. Scale bar, 200 μm. **D**, qRT-PCR analysis of expression of vascular markers genes (*Pecam1, Cld5, Cdh5, Podxl*) from tumors formed in WT and *Esm1* KO mice. n=3 replicates; *\*P=0.0101; **P=0.0141; ***P=0.0017; ****P=0.0003*. **E**, Gene Set Enrichment Analysis (GSEA) plots comparing gene expression in tumors formed in *Esm1* KO vs. WT mice. All quantitative data are average ±SD.

**Supplementary Figure 3:**
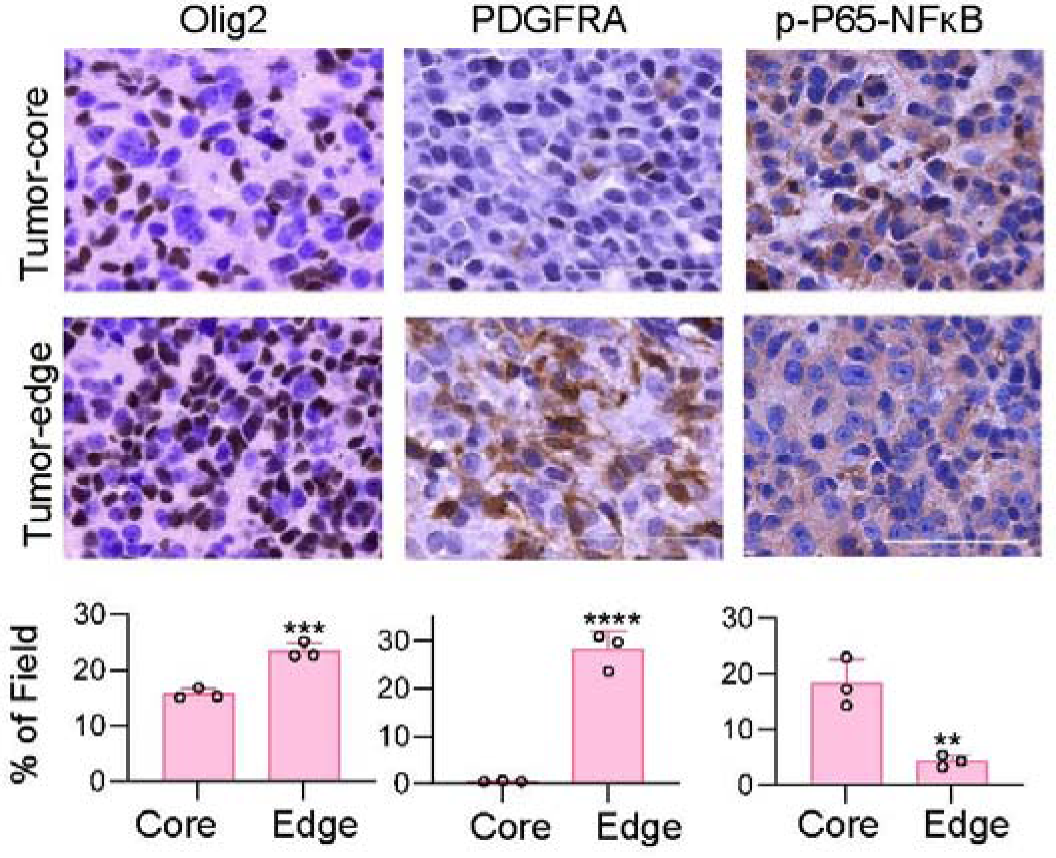
Representative IHC images of core and edge regions of the tumors obtained as in “Fig 3D” and stained for PDGFRA, Olig2 and anti-pP65-NF-κB (upper panel). % of field positive areas for the indicated proteins (lower panel). Samples from n=5 mice. *\*\*P=0.0062*, *\*\*\*P=0.0015*, *\*\*\*\*P=0. 0003.Scale* bar, 200µm. All quantitative data are average ±SD.

**Supplementary Figure 4:**
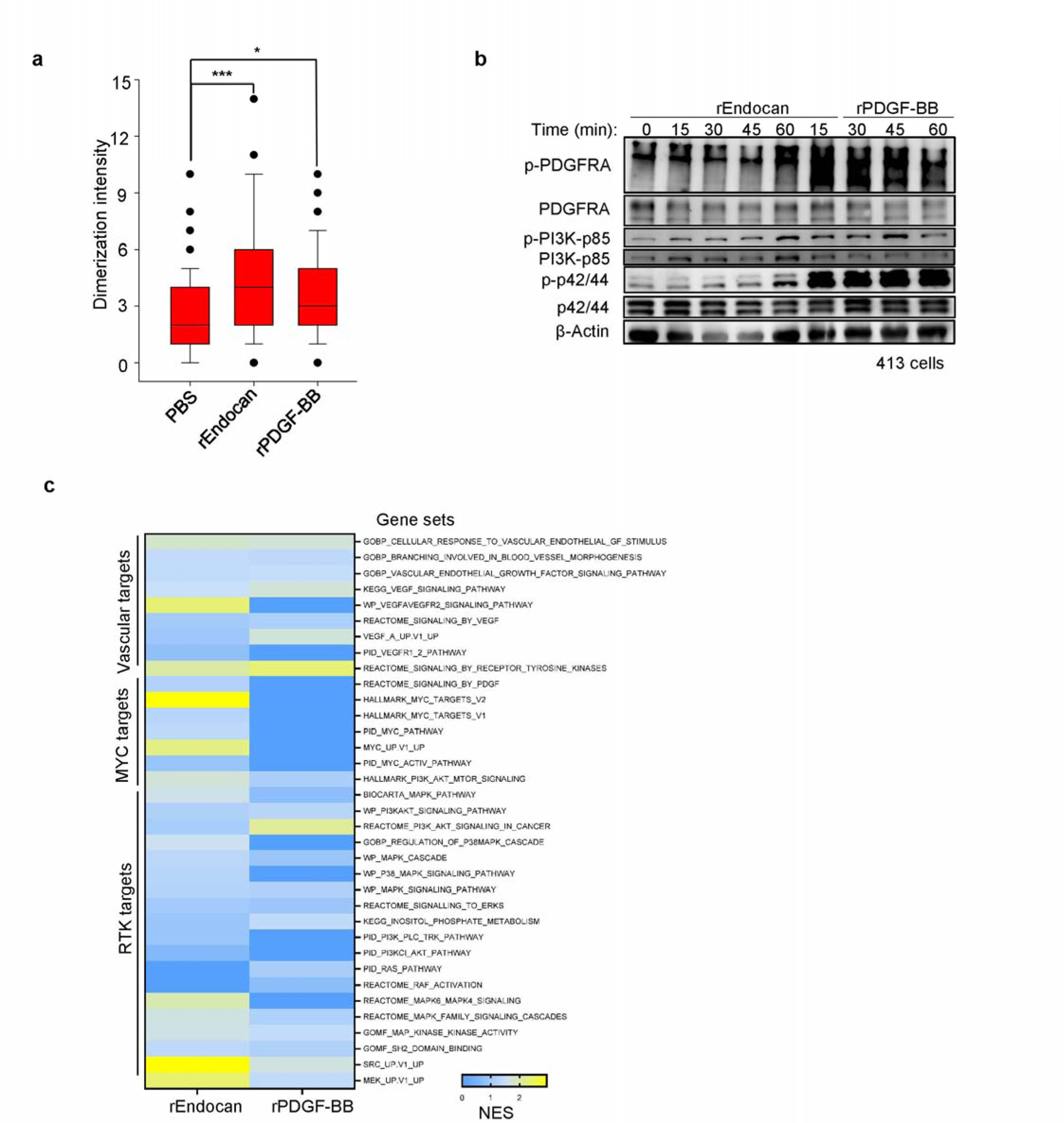
**A,** PDGFRA dimerization intensity determined by *In situ* Proximity Ligation assay (PLA) in 157 cells treated with rEndocan, rPDGF-BB or PBS (negative control). \**P*<0.05, \*\*\**P*<0.001. **B**, Western blot analysis of 413 cells incubated with 10ng/ml rEndocan or rPDGFBB for the indicated periods of time. **C,** Heatmap showing normalized enrichment scores (NES) calculated from RNAseq data of 413 cells treated with 10ng/ml rEndocan or rPDGF-BB for 48 hours, n=3 biological replicates.

**Supplementary Figure 5:**
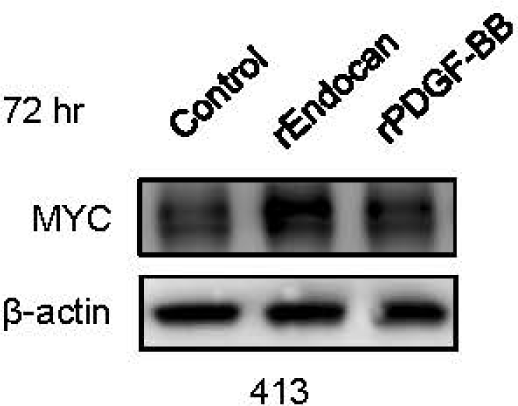
Western blot analysis of 413 cells treated with 10ng/ml rEndocan and rPDGF-BB for 72 hours.

**Supplementary Figure 6:**
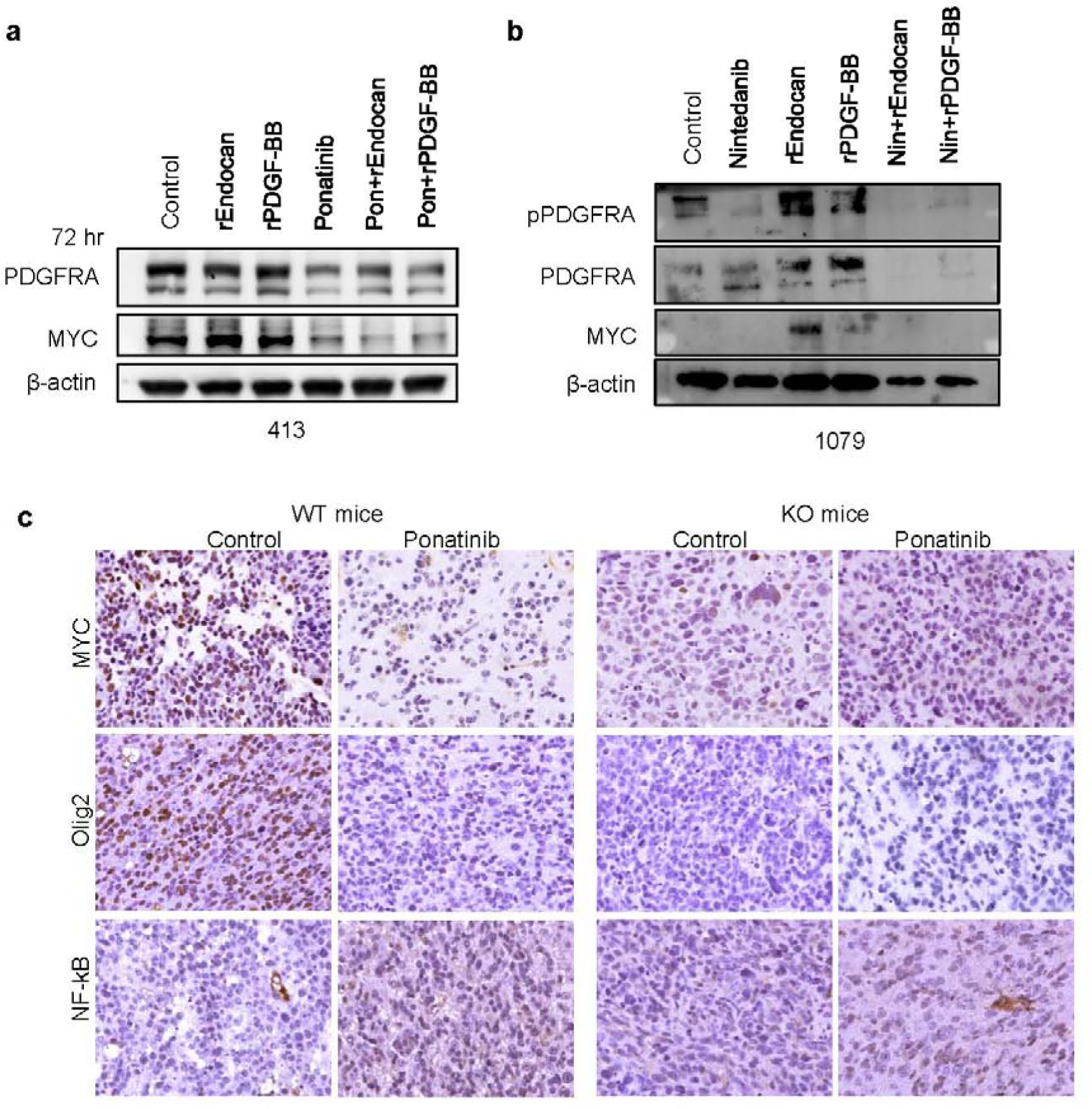
**A,** Western blot analysis of 413 cells treated with rEndocan, rPDGF-BB, ponatinib (Pon) and their combination treatment for 72 hours. **B,** Western blot analysis of 1079 cells treated with rEndocan, rPDGF-BB, Nintedanib and their combination treatment for 72 hours. **C**, Representative IHC staining for MYC, Olig2, and P65-NF-κB of tumors formed by 7080 cells in *Esm1* WT or KO mice that were treated with ponatinib or the solvent. Scale bar, 100µm.

**Supplementary Figure 7:**
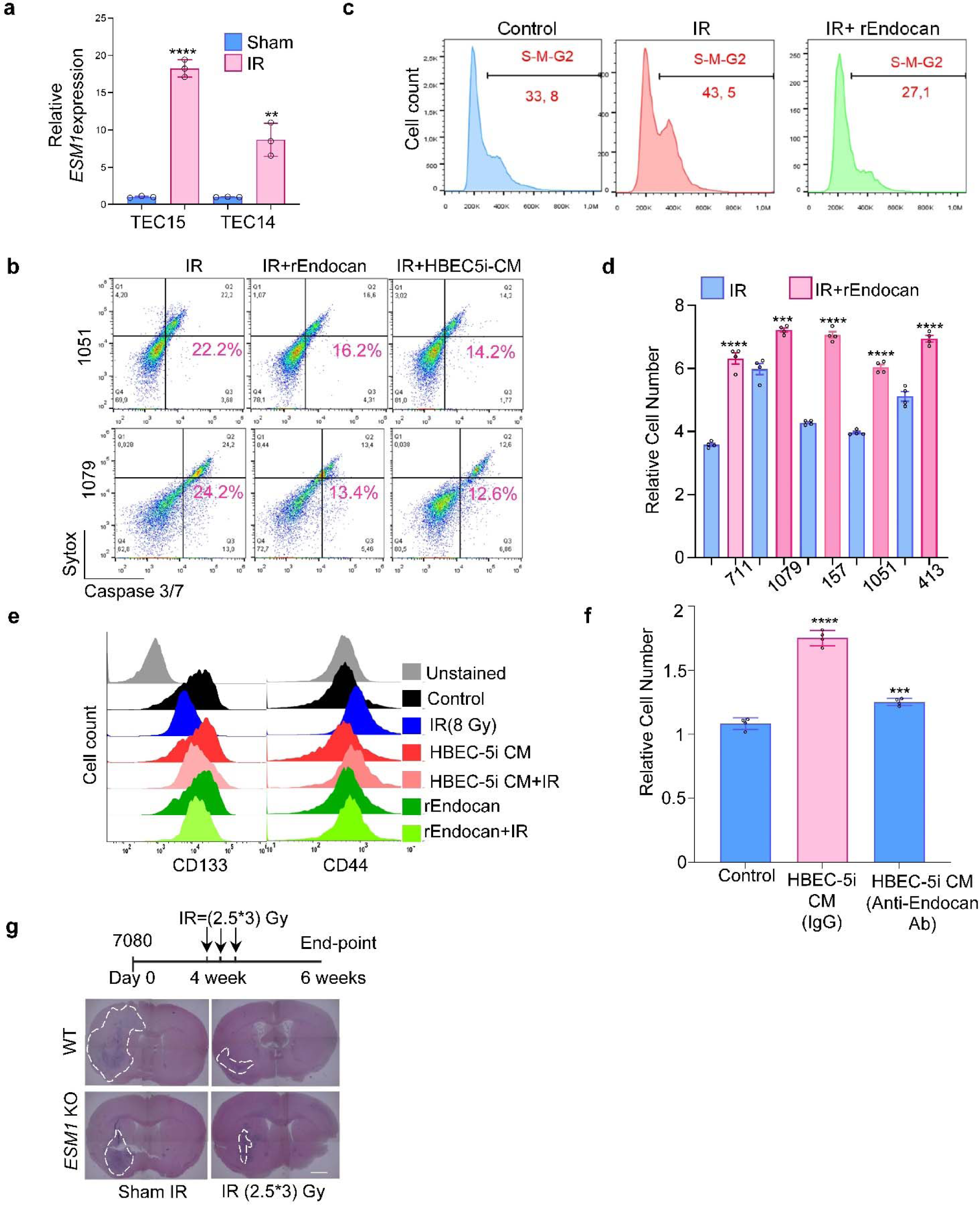
**A**, qRT-PCR analysis of *ESM1* gene expression in TEC15 and TEC14 VE cells 48 hours after treatment with 8 Gy radiation; n=3 independent experiments, *\*P=0.0039; ****P<0.0001*. **B**, FACS analysis of caspase 3/7 and SYTOX staining of 1051 cells that were preincubated with rEndocan or HBEC-5i CM for 3 days, irradiated with 8 Gy and stained two days later. **C**, FACS analysis of cell cycle distribution of 1051 cells that were left untreated, irradiated with 8Gy or preincubated for 3 days with rEndocan and subsequently irradiated. **D**, *In vitro* radio-sensitivity assay of gliomaspheres (711, 1079, 157, 1051, and 413) pretreated with rEndocan for 3 days and irradiated with 8Gy. Cell growth was measured on day 5 after irradiation. n=4 independent experiments. **E**, FACS analysis of CD133 and CD44 staining of 1079 cells pretreated with rEndocan or HBEC-5i CM for 3 days, irradiating with 8Gy and stained two days later. Unstained cells were used as control*. ***P<0.001, ****P<0.0001*. **F,** *In vitro* radio-sensitivity assay of 1079 cells preincubated for 3 days with control media, or HBEC-5i CM that was incubated with Endocan blocking antibody (Endocan depleted CM), or isotype control antibody. After incubation cells were irradiated at 8 Gy and analyzed after 3 more days. n=4 independent experiments. \*\*\**P=0.0008, ****P<0.0001*. **G**, Representative H&E images of brain sections obtained from *Esm1* WT or *Esm1* KO mice intracranially injected with 7080 cells and subsequently irradiated with 3 doses of 2.5 Gy 14 days after injection. n=5 mice per group. Scale bar, 500µm. All quantitative data are average ±SD.

